# The Cyclimids: Degron-inspired cereblon binders for targeted protein degradation

**DOI:** 10.1101/2023.04.22.537935

**Authors:** Saki Ichikawa, N. Connor Payne, Wenqing Xu, Chia-Fu Chang, Nandini Vallavoju, Spencer Frome, Hope A. Flaxman, Ralph Mazitschek, Christina M. Woo

## Abstract

Cereblon (CRBN) is an E3 ligase substrate adapter widely exploited for targeted protein degradation (TPD) strategies. However, achieving efficient and selective target degradation is a preeminent challenge with ligands that engage CRBN. Here, we report that the cyclimids, ligands derived from the C-terminal cyclic imide degrons of CRBN, exhibit distinct modes of interaction with CRBN and offer a facile approach for developing potent and selective bifunctional degraders. Quantitative TR-FRET-based characterization of 60 cyclimid degraders in binary and ternary complexes across different substrates revealed that ternary complex binding affinities correlated strongly with cellular degradation efficiency. Our studies establish the unique properties of the cyclimids as versatile warheads in TPD and a systematic biochemical approach for quantifying ternary complex formation to predict their cellular degradation activity, which together will accelerate the development of degraders that engage CRBN.

## Main Text

Targeted protein degradation (TPD) induced by small molecules is emerging as an attractive strategy for eliminating disease-relevant proteins from cells.^1-3^ Through chemically induced proximity, small molecule degraders, such as molecular glues^4^ and proteolysis targeting chimeras (PROTACs),^1-3,5^ recruit target proteins to an E3 ligase complex and promote their subsequent ubiquitination and proteasomal degradation. Among the >600 E3 ligases encoded by the human genome, cereblon (CRBN), a substrate adapter in the CRL4^CRBN^ E3 ligase complex,^6-8^ is one of the best-characterized substrate adapters for TPD applications. CRBN was originally discovered in association with intellectual disability^9^ and was later identified as the primary target of thalidomide.^6^ Thalidomide and its derivatives, lenalidomide and pomalidomide, are members of a class of pharmaceutical agents termed the immunomodulatory drugs (IMiDs); the IMiDs have unique and pleiotropic clinical properties against multiple myeloma,^10,11^ del(5q) myelodysplastic syndrome,^12^ and other hemopoietic malignancies,^13^ as well as anti-inflammatory effects in the treatment of inflammatory diseases including erythema nodosum leprosum (ENL), rheumatoid arthritis, HIV-associated aphthous ulcers, and tuberculous meningitis.^14^ These effects are partially realized through binding to CRBN, thereby altering its substrate selection (*e.g.,* IKZF1, IKZF3, CK1α, or GSPT1) for ubiquitination and degradation.^10-13^ Engagement of CRBN using heterobifunctional degraders (*e.g.*, PROTACs) by linking an IMiD to a ligand for a target protein has also spurred the development of many IMiD-based PROTACs for degradation of a myriad of target proteins and is a promising new modality in both chemical biology and drug discovery.^15-18^

Despite the immense promise of degrader modalities, challenges in the development of efficient degrader molecules include (i) achieving high target selectivity, (ii) identifying and parsing through the diverse modes of target–degrader–E3 ligase ternary complexes, (iii) engineering precise control of the ternary complex orientation, and (iv) optimizing cellular degradation activity and characterizing these molecules in biochemical and cellular assays.^17^ Heterobifunctional degraders consist of two ligands, one for interacting with a target protein and the other for recruiting an E3 ligase, tethered by a linker. The design of heterobifunctional degraders through linker screening (termed “linkerology”)^17,19^ offers one approach to tackle challenges in degrader discovery and optimization. The linker controls critical aspects of degrader design, such as the solution conformation of the degrader molecule,^20^ as well as ternary complex engagement^21^ and orientation,^22^ thereby influencing target selectivity and degradation efficiency. As the formation of a productive ternary complex is crucial for efficient degradation events, many biophysical approaches are available to measure PROTAC-mediated ternary complexes, such as proximity-based assays (*e.g.*, AlphaScreen/AlphaLISA),^23,24^ isothermal titration calorimetry,^25^ surface plasmon resonance,^26^ and other cell-based assays (*e.g.*, separation of phase-based protein interaction reporter [SPPIER]^27^ and bioluminescence resonance energy transfer [BRET]^28^). While each assay is associated with different strengths and weaknesses, a unified, quantitative, and ideally high-throughput approach to derive a systematic correlation between the ternary complex formation and degradation efficacy of a heterobifunctional degrader would greatly accelerate the development of these novel modalities.

A mechanism to systematically evaluate ternary complex formation by variation of the CRBN warhead, rather than the linker, may provide an orthogonal and tunable strategy to generate productive ternary complexes for TPD. Indeed, minor structural changes on the CRBN ligand can dramatically alter biological activities in cells,^22^ and the engagement of CRBN tends to be strongest with a 6-membered glutarimide in the IMiD structure and is significantly diminished when substituted with a 7-membered^29^ or 5-membered^30^ cyclic imide. Recently, we discovered that C-terminal cyclic imides are previously overlooked post-translational modifications that arise from intramolecular cyclization of glutamine or asparagine, which afford a degron for the thalidomide-binding domain of CRBN (Fig. 1a).^31^ CRBN flexibly recognizes a range of C-terminal cyclic imide dipeptides when conjugated to JQ1 to afford functional degraders for BRD4 (bromodomain-containing protein 4)^32^ with intriguing differences in degradation efficiency that may be attributed to one of several factors, including differential engagement of CRBN, positive or negative ternary complex formation, or cellular permeability. Notably, both 6-membered cyclic imides and 5-membered cyclic imides served as effective degrons for CRBN, implying that the properties of these degron motifs may be in stark contrast to those of the IMiDs when embedded in a bifunctional degrader and thus render new chemical space accessible. In this study, we designed and evaluated several classes of PROTACs bearing cyclic imide ligands, termed cyclimids, against a series of cellular targets. Employing a quantitative, high-throughput TR-FRET-based ternary complex assay platform, our results provide evidence that the cyclimids induce distinct modes of ternary complex formation between CRBN and target proteins in a linker-independent manner. The cyclimids show higher on-target selectivity via minimization of off-target CRBN-mediated molecular glue activity and maximization of ternary complex potency. Overall, our results demonstrate the potential for the cyclimids to yield potent and efficient PROTACs through systematic tuning of target–CRBN ternary complexes.

**Fig. 1.**
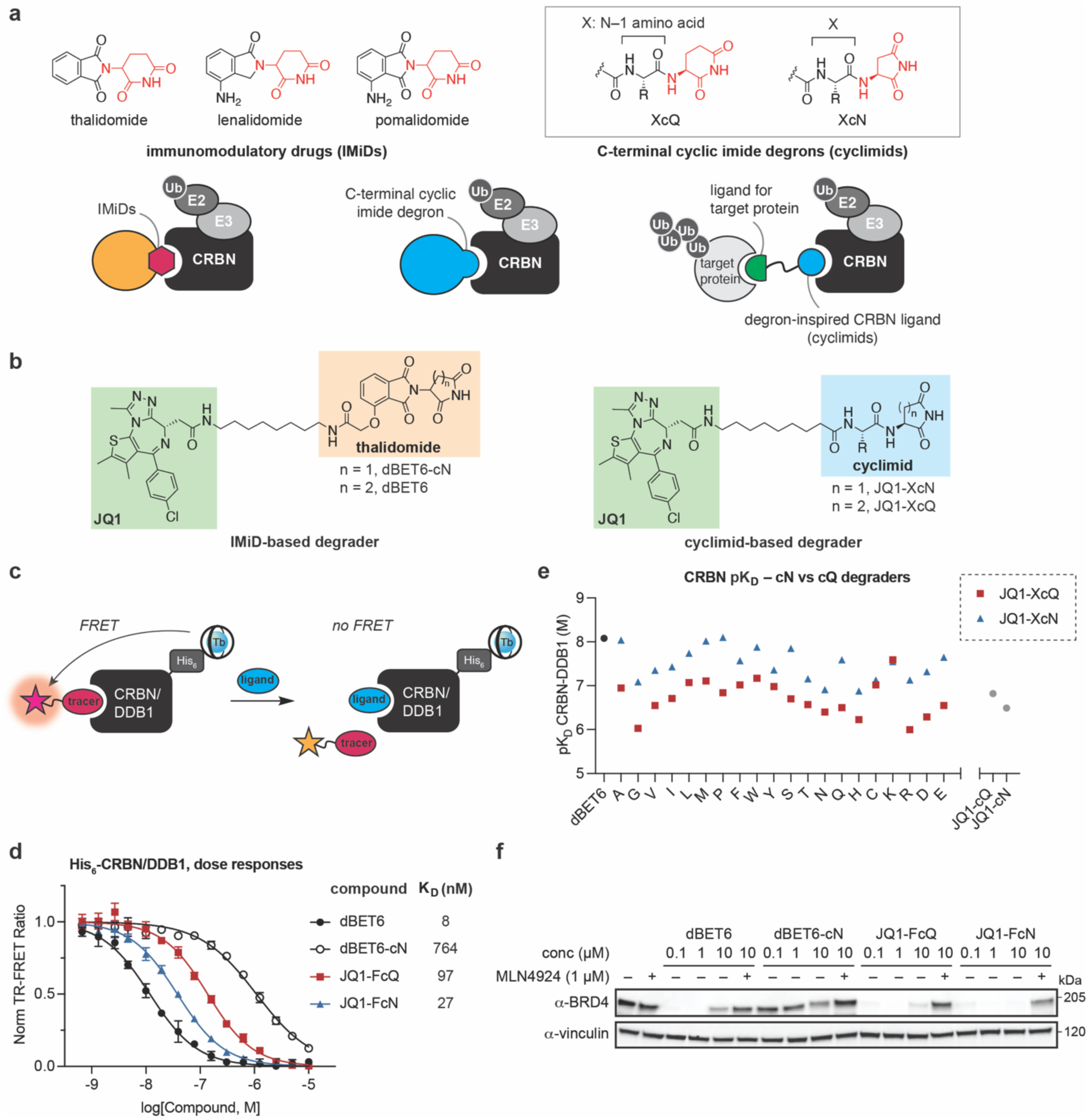
The Cyclimids, degron-inspired CRBN ligands, exhibit distinct and diverse binding affinities against CRBN compared to IMiDs. (**a**) CRBN recognizes the C-terminal cyclic imide degrons, which are mimicked by thalidomide, lenalidomide, and pomalidomide. Structures of the immunomodulatory drugs (IMiDs) and the C-terminal cyclic imide degrons (XcQ and XcN, collectively called the cyclimids). The Cyclimids, a class of CRBN ligands inspired by the C-terminal cyclic imide degron of CRBN, may serve as unique CRBN warheads to develop heterobifunctional degraders. (**b**) Structures of IMiD-based degraders (dBET6 and dBET6-cN) and cyclimid-based degraders (JQ1-XcQ and JQ1-XcN) for functional engagement of CRBN and BRD4 degradation in cells. (**c**) Schematic of the TR-FRET ligand-displacement assay for His_6_-CRBN/DDB1. (**d**) Dose-titration of the indicated degraders in TR-FRET ligand displacement assays with His_6_-CRBN/DDB1 and the determined equilibrium dissociation constants (K_D_) of the indicated compounds against the His_6_-CRBN/DDB1 complex. (**e**) Comparison between the pK_D_ values of cN/cQ cyclimid- and IMiD-based degraders against CRBN/DDB1. In contrast to IMiD-based degraders, cN-cyclimid-based degraders generally bind to CRBN/DDB1 more tightly than their cQ counterparts. Data in (**d**) and (**e**) are presented as mean ± SD (n = 3 technical replicates). (**f**) Western blot of BRD4 after treatment of HEK293T cells for 4 h with dBET6, dBET6-cN, JQ1-FcQ, or JQ1-FcN over a 100 nM–10 μM dose-response range. Western blot data are representative of three independent replicates. For uncropped western blot images, see Supplementary Figure 2.

### The cyclimids exhibit distinct and diverse binding affinities against CRBN

To characterize the cyclimids as a unique class of CRBN ligands for TPD applications, we first constructed a library of 40 cyclimid-based degraders against BRD4. The cyclimid-based BRD4 degrader library is composed of a series of twenty 6-membered cyclic glutarimide (cQ)-bearing cyclimid BRD4 degraders (JQ1-XcQ),^31^ and an additional set of twenty cyclimid degraders consisting of JQ1 linked to a dipeptide with a 5-membered cyclic aspartimide (cN) (JQ1-XcN; Fig. 1b). We installed all 20 canonical amino acids at the N–1 position (X) to the cyclic imide motif (cQ or cN) when constructing the library of cyclimids (XcQ or XcN). For direct comparison, we used the IMiD-based degrader dBET6^15,16,22^ and prepared its cN counterpart with no amino acid at the N–1 position (dBET6-cN; Fig. 1b). The same aliphatic linker was used across all cyclimid degraders to control for its contribution to degrader properties.

We first assessed the binding of the cyclimid-based degraders to CRBN by a time-resolved Förster resonance energy transfer (TR-FRET) ligand displacement assay using Thal-FITC as a tracer molecule, specific for the thalidomide-binding domain of CRBN (Fig. 1c, Extended Data Fig. 1a). Saturation binding of Thal-FITC against full-length His_6_-CRBN/DDB1 detected by a CoraFluor-1-labeled anti-His_6_ antibody^33^ yielded an equilibrium dissociation constant (K_D_) of 117 nM for His_6_-CRBN/DDB1, which is comparable to the K_D_ values of thalidomide measured by fluorescence polarization (Extended Data Fig. 1b).^34^ We next determined the binding affinities of the 40 cyclimid-based degraders, along with the IMiD-based degraders dBET6 and dBET6-cN, against CRBN/DDB1 by competition binding (Fig. 1d–e). The cyclimid degraders, JQ1-FcQ and JQ1-FcN, and dBET6 demonstrated similar levels of binding to CRBN (K_D_ values < 100 nM), whereas dBET6-cN exhibited more limited engagement of CRBN (K_D_ = 764 nM, Fig. 1d, **Supplementary Table 1**). All 40 cyclimid degraders functionally engaged CRBN and displayed K_D_ values ranging from 8 nM–1 μM, depending on the amino acid installed at the N–1 position (Fig. 1e, **Supplementary Table 1**). The degraders JQ1-cQ and JQ1-cN, which lack the amino acid at the N–1 position, also engaged CRBN at a moderate level independent of the ring size of the cyclic imide (Fig. 1e, Extended Data Fig. 1c). These data suggest that the additional amino acid at the N–1 position can tune binding affinities against CRBN in the cyclimid degraders. Notably, the cyclimid degraders with 5-membered cN generally exhibit higher affinities to CRBN than their 6-membered cQ counterparts (Fig. 1e).

The selected monofunctional cyclimids were also evaluated as ligands for CRBN, independent of the linker to JQ1 (Extended Data Fig. 1d–f). We observed a similar trend in which the binding affinity towards CRBN with cN-bearing dipeptides was generally superior to that of its cQ counterpart (Extended Data Fig. 1d). In contrast, pomalidomide (POM) exhibited much higher binding affinity than the 5-membered analog POM-cN^30^ (∼500-fold difference in K_D_, Extended Data Fig. 1f). The inferior affinity of POM-cN compared to the parent cQ counterpart POM may be related to its inflexible projection of the isoindoline-1,3-dione backbone caused by the smaller ring size, which prevents favorable interactions with nearby residues at the thalidomide-binding domain of CRBN (Extended Data Fig. 1g–h). To further investigate the functional difference between 5-membered cN embedded in IMiD or cyclimid scaffolds, the BRD4 degradation activity of dBET6-cN and JQ1-FcN, along with 6-membered cQ-based degraders dBET6 and JQ1-FcQ, were assessed in HEK293T cells (Fig. 1f). While dBET6 and JQ1-FcQ exhibited similar levels of BRD4 degradation, JQ1-FcN demonstrated efficient degradation of BRD4 at a concentration of 100 nM, whereas dBET6-cN was ineffective even at a concentration of 10 μM. The degradation efficiency directly correlates with the difference in binding affinity of cN against CRBN when incorporated within either IMiD or cyclimid degraders.

### Ternary complexes induced by cyclimid degraders are tunable and distinct from IMiD-based degraders

In addition to the distinct and diverse binding affinities of the cyclimids toward CRBN, we hypothesized that these cyclimid degraders may induce differential ternary complex modes relative to the IMiD-based degraders and, therefore, screening the amino acid at the N–1 position, installed next to the cyclic imide motif, may afford rapid access to productive ternary complexes. We thus sought to evaluate the ternary complex formation between CRBN and each bromodomain of the target protein BRD4 (BD1 and BD2) induced by our complete panel of cyclimid or IMiD-based degraders. To circumvent the difficulty of extracting the absolute values of binding affinities for a three-body binding system,^35^ we exploited the fact that a complex three-component system under certain conditions can be rendered comprehensible by applying familiar concepts from simpler binary complex equilibria, as conceptually demonstrated by multiple groups using biophysical assays, such as isothermal calorimetry (ITC), surface plasmon resonance (SPR), biolayer interferometry (BLI), and fluorescence polarization (FP).^25,26,35-37^ However, these techniques suffer from either low-throughput, high material requirements, or low sensitivity, and therefore are often only used to characterize the most promising PROTAC candidates in a given medicinal chemistry platform. Therefore, a quantitative, high-throughput, and straightforward assay to biochemically evaluate PROTAC-mediated ternary complexes would be highly desirable.

To this end, we sought to modify our Thal-FITC/His_6_-CRBN/DDB1 TR-FRET ligand displacement platform to enable quantitative ternary complex evaluation. Specifically, when [BRD4] ≫ K_D, BRD4_ and K_D, CRBN_ ≫ [CRBN], our TR-FRET ligand displacement platform can instead provide the equilibrium dissociation constant of the degrader–target binary complex for CRBN (K_D, CRBN_[ternary], Fig. 2a). Saturation of the BRD4-targeting ligand (*i.e.,* JQ1) can be achieved by using a high concentration of recombinant BD1 or BD2 in the assay solution, resulting in the formation of a binary complex which can compete with Thal-FITC in the TR-FRET ligand displacement assay against CRBN. To achieve near-saturation, the minimum concentration of GST-tagged BRD4 bromodomains BD1 or BD2 was first determined by the TR-FRET ligand displacement assays using JQ1-FITC^38^ as a tracer (Extended Data Fig. 2, **Supplementary Table 1**). All BRD4-targeting cyclimid degraders possess K_D_ values for BRD4(BD1) and BRD4(BD2) in the 20–100 nM range. As such, at a concentration of 1 μM BRD4(BD1) or BRD4(BD2) (approximately ∼10–50-fold greater than the K_D_ value), this will result in essentially quantitative formation of cyclimid–BD1/2 binary complexes (corresponding to 92–98% formation of binary complex^39^). Furthermore, the cooperativity of binding can be directly calculated through these assays, which gives the cooperativity factor Alpha as a quantitative and comparable indicator to evaluate the ternary complex formation.^22,35^ The cooperativity factor Alpha is defined by:

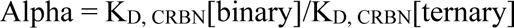

**Fig. 2.**
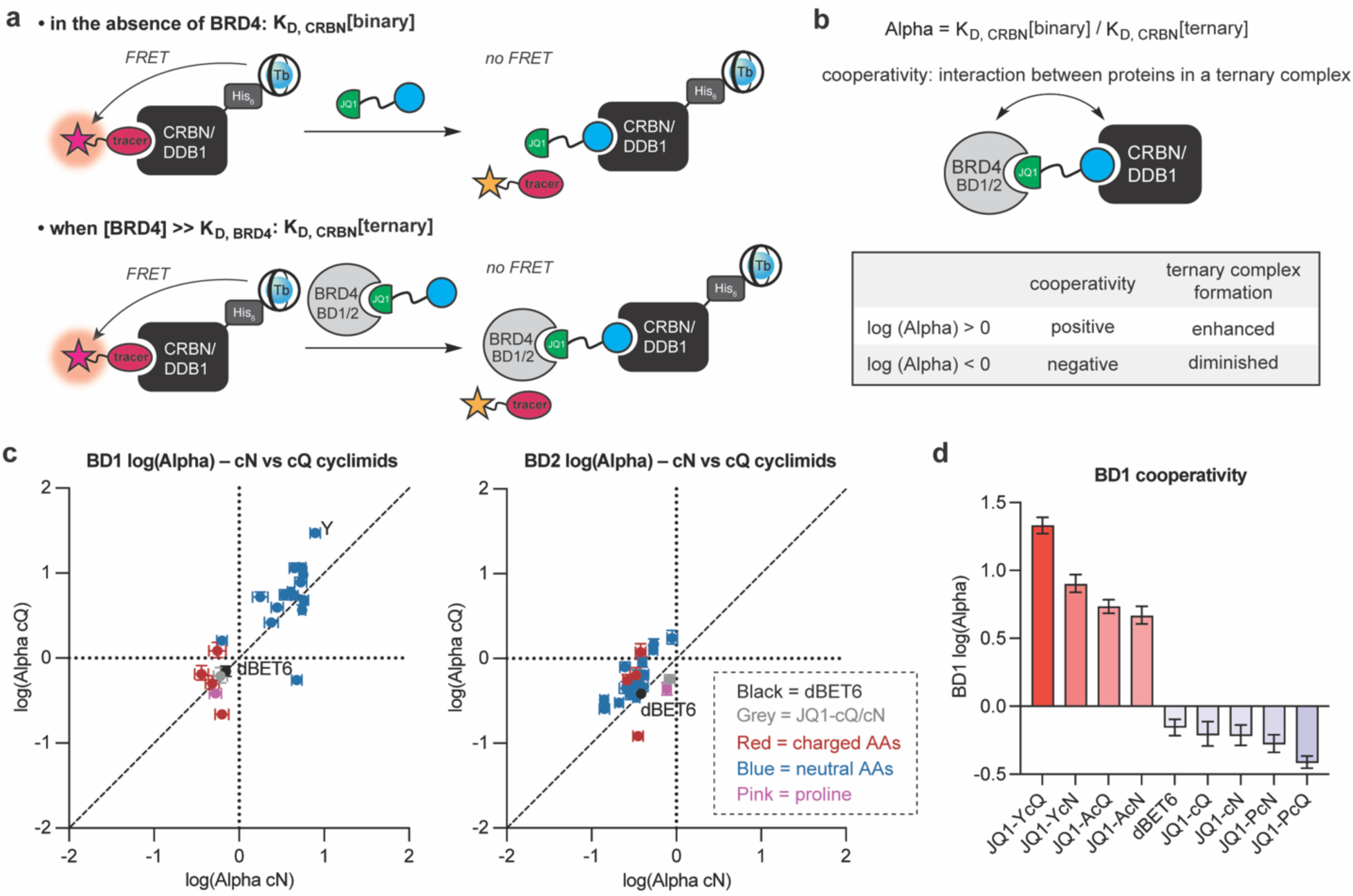
Cyclimid degraders induce ternary complexes that are tunable and distinct from IMiD-based degraders. (**a**) Schematic of the TR-FRET assay principle for determining K_D_[binary] and K_D_[ternary] against CRBN/DDB1. K_D, CRBN_[ternary] values of the compounds were measured using the TR-FRET ligand displacement assay in the presence of a near-saturating concentration of either BD1 or BD2 domain of BRD4 (1 µM). (**b**) Schematic of the cooperativity factor Alpha. (**c**) Comparison of log(Alpha) values between cN and cQ cyclimid degraders. Log(Alpha) values for the cyclimids with the BD1 domain of BRD4 showed significant variance depending on the amino acid at the N–1 position. Data are presented as mean ± SD (n = 3 technical replicates). (**d**) Log(Alpha) for the selected cyclimid degraders with the BD1 domain of BRD4. Error bars represent mean ± SD.

where a positive cooperativity between two proteins results in log(Alpha) > 0, which suggests the direct interprotein interaction upon the ternary complex formation is favorable, while a negative cooperativity [log(Alpha) < 0] disfavors interprotein contacts (Fig. 2b). Because JQ1 binds to both BD1 and BD2 bromodomains with comparable affinities (K_D_ = 50 nM for BD1 and 90 nM for BD2),^32^ we focused on whether the structurally diverse cyclimid-based degraders would interact with each bromodomain differently upon formation of a ternary complex with CRBN.

Using this assay design, we measured K_D, CRBN_[ternary, BD1 or BD2] and calculated the cooperativity factor Alpha for all 40 cyclimid degraders between His_6_-CRBN/DDB1 and GST-BD1 or BD2 (**Supplementary Table 1**). The cyclimid degraders exhibited diverse cooperativity profiles compared to the IMiD-based degrader dBET6, and the cooperativity of binding was dramatically altered by the amino acid installed at the N–1 position (Fig. 2c). The cyclimid degraders with neutral amino acids displayed positive cooperativity with BRD4(BD1) and negative cooperativity with BRD4(BD2), while the IMiD-based degrader dBET6 exhibited negative cooperativity for both bromodomains of BRD4, consistent with the previous report.^22^ Additionally, degraders without the amino acid at the N–1 position, JQ1-cQ and JQ1-cN, showed no significant cooperativity for either bromodomain, providing evidence that the N–1 amino acid is responsible for driving the differences in ternary complex formation. The log(Alpha) values of the cyclimid degraders varied quite significantly (*e.g*., an 80-fold difference in Alpha for BD1 between JQ1-YcQ and JQ1-PcQ; Fig. 2d, **Supplementary Table 1**). To further assess the selectivity of ternary complex formation mediated by the cyclimids with BD1 versus BD2 domains of BRD4, we calculated the BD1 and BD2 selectivity parameters derived from Alpha(BD1)/Alpha(BD2) for each degrader (Extended Data Fig. 3a). Among all cyclimid degraders tested, JQ1-YcQ showed the greatest selectivity for the ternary complex formation with BRD4(BD1), with a 57-fold difference in Alpha for BD1 and BD2 (Extended Data Fig. 3b, **Supplementary Table 1**). Collectively, these data demonstrate that the key to the diversity of ternary complex structures formed by the cyclimids, as evidenced by K_D, CRBN_[ternary] value and cooperativity factor Alpha, lies in the amino acid at the N–1 position adjacent to the cyclic imide motif, especially considering they all possess the same linker motif.

**Fig. 3.**
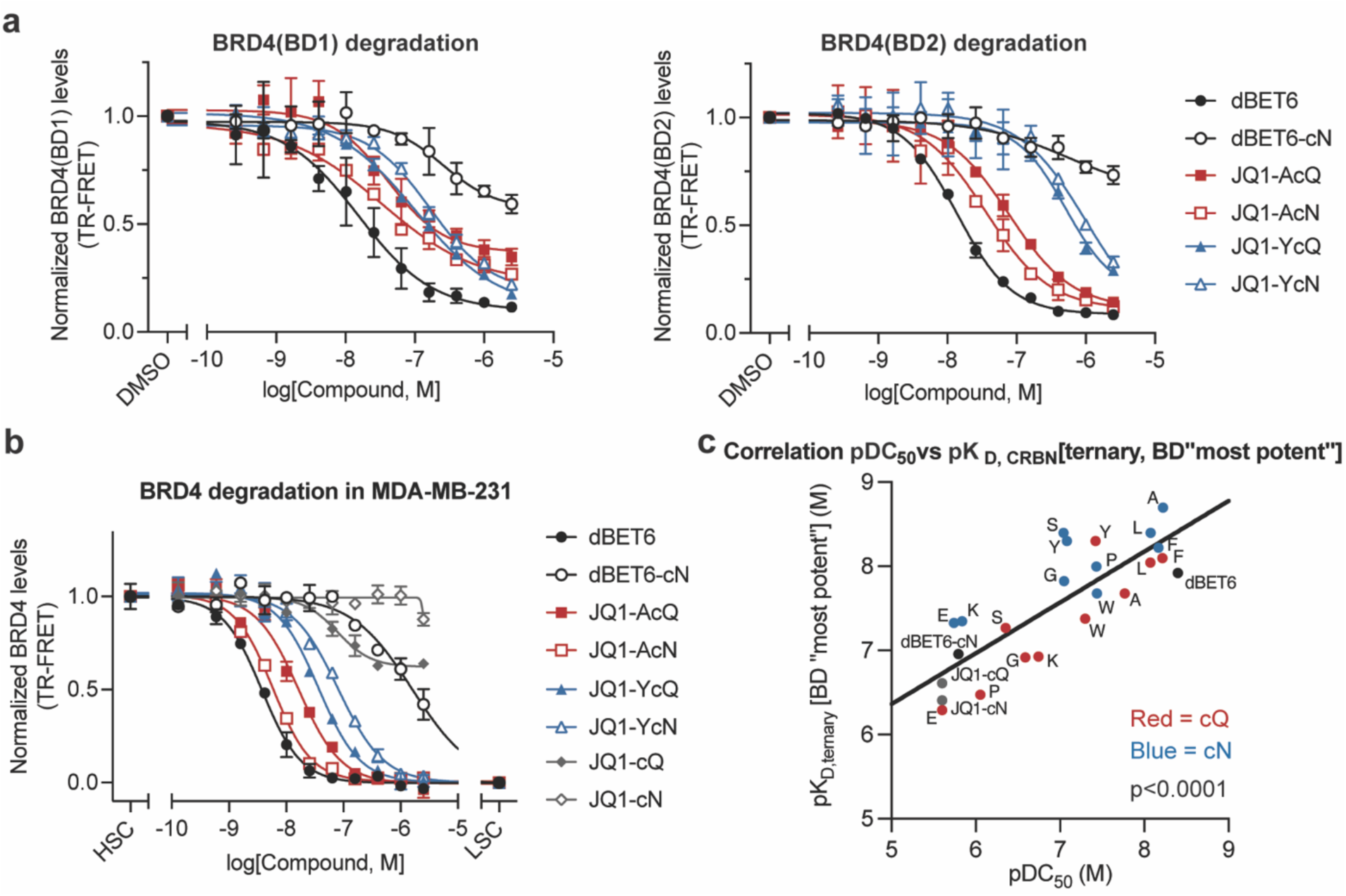
K_D, CRBN_[ternary] values are predictive of the cellular degradation activity of cyclimid degraders. (**a**) BRD4(BD1)-GFP or BRD4(BD2)-GFP levels in HEK293T cells stably expressing BRD4(BD1)-GFP or BRD4(BD2)-GFP after 4 h treatment with dBET6 and the selected cyclimid BRD4 degraders were measured by TR-FRET assay. For assay design, see Extended Data Fig. 4a. (**b**) Endogenous BRD4 protein levels in MDA-MB-231 cell lysate after 5 h treatment with dBET6 and the selected cyclimid BRD4 degraders were measured by TR-FRET assay. HSC, high signal control; LSC, low signal control. Data in (**a**) and (**b**) are presented as mean ± SD (n = 2 biologically independent samples). (**c**) Correlation between pDC_50_ values of the selected cyclimid degraders for endogenous BRD4 degradation and their biochemical pK_D, CRBN_[ternary] values. In the correlation graph, for each degrader, the most potent pK_D, CRBN_[ternary] value between pK_D, CRBN_[ternary, BD1] and pK_D, CRBN_[ternary, BD2] was selected.

### K_D, CRBN_[ternary] values are predictive of the cellular degradation activity of cyclimid degraders

In light of the significance of the amino acid at the N–1 position upon ternary complex formation, we next asked whether the ternary complex induced by the cyclimid degraders would be predictive of the cellular degradation activity of these degraders. Prior art indicates a strong correlation between cooperativity, ternary complex K_D_ or half-lives, and the degradation potency of the heterobifunctional degrader, although this trend has been qualitatively demonstrated with select lead compounds across multiple reports.^23,25,26,36,40-45^ We therefore sought to extend these important precedents by conducting absolute quantitative measurements of the binary affinity, ternary affinity, and degradation on a scale that has not been previously performed. Collectively, we now provide a single report detailing the thorough correlation of biochemical and cellular activity for our library of 40 cyclimid-based BRD4 degraders and extend these observations to an additional 20 cyclimid degraders for additional targets.

First, to assess the selectivity in the degradation of each bromodomain of BRD4, we selected two pairs of cyclimid degraders, JQ1-YcQ/YcN and JQ1-AcQ/AcN, and the IMiD-based analogs dBET6/dBET6-cN for evaluation in reporter HEK293T cells stably expressing either BRD4(BD1)-GFP or BRD4(BD2)-GFP.^22^ These two pairs of the cyclimid degraders were selected because JQ1-AcQ/AcN showed little to no ternary complex selectivity in our biochemical ternary complex assay, whereas JQ1-YcQ/YcN displayed 57- and 27-fold biochemical selectivity for ternary complexes with BD1. The induced degradation of BRD4(BD1) or BRD4(BD2) in these reporter cells after 4 h incubation with the selected cyclimid degraders or the IMiD-based degraders was quantified directly in lysates adapting a TR-FRET-based strategy utilizing CoraFluor-labeled anti-GFP nanobodies^46^ (Fig. 3a, Extended Data Fig. 4a–b, **Supplementary Table 1**). Both JQ1-YcQ and JQ1-YcN exhibited higher potency for BD1 degradation with a ∼5-10-fold difference in half-maximal degradation concentrations (DC_50_) between BD1 and BD2, in good agreement with the BD1/BD2 selectivity parameter predicted from ternary complex studies [JQ1-YcQ: DC_50_(BD1) = 103 nM; DC_50_(BD2) = 513 nM, JQ1-YcN: DC_50_(BD1) = 168 nM; DC_50_(BD2) = 1.01 µM, **Supplementary Table 1**]. In contrast, the other pair of cyclimid degraders, JQ1-AcQ and JQ1-AcN, as well as dBET6, displayed relatively equal activity toward both bromodomains [JQ1-AcQ: DC_50_(BD1) = 41 nM; DC_50_(BD2) = 78 nM, JQ1-AcN: DC_50_(BD1) = 34 nM; DC_50_(BD2) = 36 nM, dBET6: DC_50_(BD1) = 15 nM; DC_50_(BD2) = 14 nM, **Supplementary Table 1**], consistent with a lack of selectivity between the individual recombinant bromodomains measured by *in vitro* ternary complex assays (Extended Data Fig. 3a). Additionally, a significant correlation was observed between pDC_50_ values for degradation of each bromodomain and pK_D, CRBN_[ternary, BD1 or BD2] values of the degraders tested (p-value=0.016, Extended Data Fig. 4c).

**Fig. 4.**
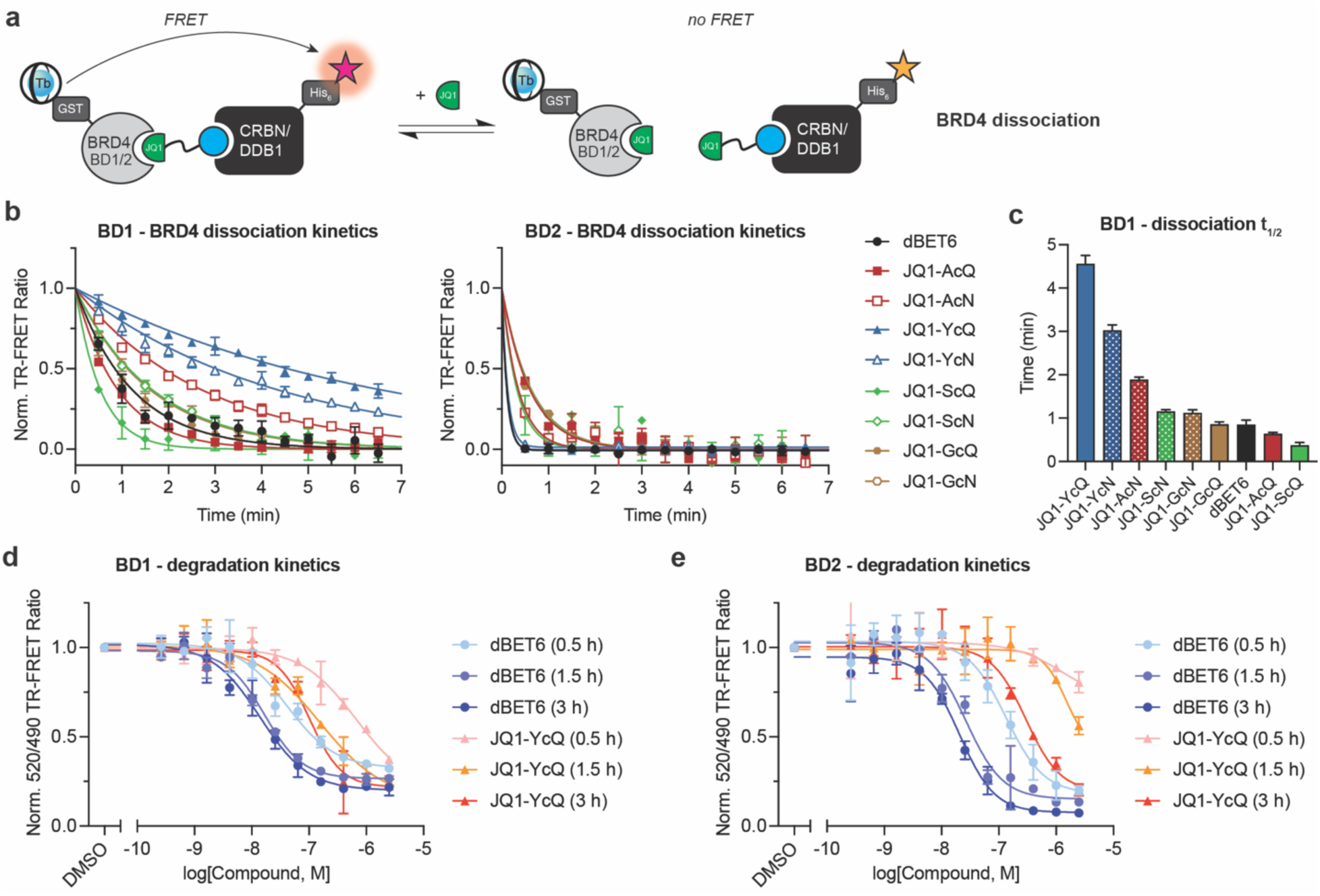
Ligand-dependent dynamics of ternary complex dissociation and degradation with the cyclimids. (**a**) Schematic illustration of TR-FRET assay system employed to monitor BRD4 dissociation from the degrader-mediated ternary complex. (**b**) Kinetic measurements of BRD4 dissociation in the presence of the BD1 or BD2 domain of BRD4 from the ternary complex induced by the selected cyclimid degraders or dBET6. Data are presented as mean ± SD (n = 2 technical replicates) and fitted to a one-phase decay model in Prism 9. (**c**) Half-lives (*t*_1/2_) of the CRBN/DDB1–BRD4(BD1) ternary complex mediated by the indicated cyclimid degraders and dBET6. Error bars represent mean ± SD (n = 2 technical replicates). (**d**) Kinetics of depletion of BRD4(BD1)-GFP in HEK293T cells stably expressing BRD4(BD1)-GFP after the treatment with JQ1-YcQ or dBET6. (**e**) Kinetics of depletion of BRD4(BD2)-GFP in HEK293T cells stably expressing BRD4(BD2)-GFP after the treatment with JQ1-YcQ or dBET6. Data in (**d**) and (**e**) are presented as mean ± SD (n = 2 biologically independent samples).

Encouraged by this observation, we next investigated the correlation between the degradation potency of endogenous BRD4 and K_D, CRBN_[ternary] values, with the intent of characterizing the K_D, CRBN_[ternary] values as a quantitative indicator for predicting the degradation potency profiles of the cyclimid degraders. To examine the degradation efficiencies of the cyclimid degraders for endogenous BRD4, MDA-MB-231 cells were treated for 5 h with a series of cyclimid degraders and controls in dose-response, followed by the quantification of endogenous BRD4 levels by TR-FRET^38^ (Fig. 3b, Extended Data Fig. 5a). This subset of the cyclimid degraders (24 of 44 degraders) was chosen to reflect nonpolar aliphatic, aromatic, and polar amino acids at the N–1 position. The cyclimid degraders exhibited a 500-fold range of DC_50_ values. The N–1 position of the cyclimids was essential for tuning the degradation properties, where in general cyclimid degraders with nonpolar or aromatic amino acid side chains at the N–1 position facilitated BRD4 degradation to a greater extent (**Supplementary Table 1**). This differential activity may be further amplified by decreased cellular permeability of analogs with polar side chains. The cellular degradation profile of the cyclimid degraders showed the strongest correlation with the pK_D, CRBN_[ternary] values against their respective “most potent” BRD4(BD1) or BRD4(BD2) domain (p-value<0.0001, Fig. 3c). Cellular degradation also correlated significantly with binary CRBN target engagement (p-value=0.007, Extended Data Fig. 5b). In contrast, no correlation was observed with target engagement toward BRD4(BD1) and BRD4(BD2) (Extended Data Fig. 5c– d), most likely because all degraders possess the same BRD4 targeting ligand (JQ1). Further evaluation of the 20 cN-bearing cyclimid degraders in HEK293T cells showed dose-dependent degradation of BRD4 equivalent to or better than that observed in MDA-MB-231 cells (Extended Data Fig. 5e–h). The cN-based cyclimid degraders generally possess superior degradation abilities compared to their cQ counterparts, in alignment with the observed differences in binding affinities to CRBN both in binary and ternary systems (**Supplementary Table 1**). In sum, these results collected from a comprehensive library of cyclimid degraders demonstrate that the degradation efficacy of cyclimid degraders can be modulated through the formation of diverse ternary complex structures, enabling K_D, CRBN_[ternary] values to serve as a quantitative indicator to predict the degradation activity.

**Fig. 5.**
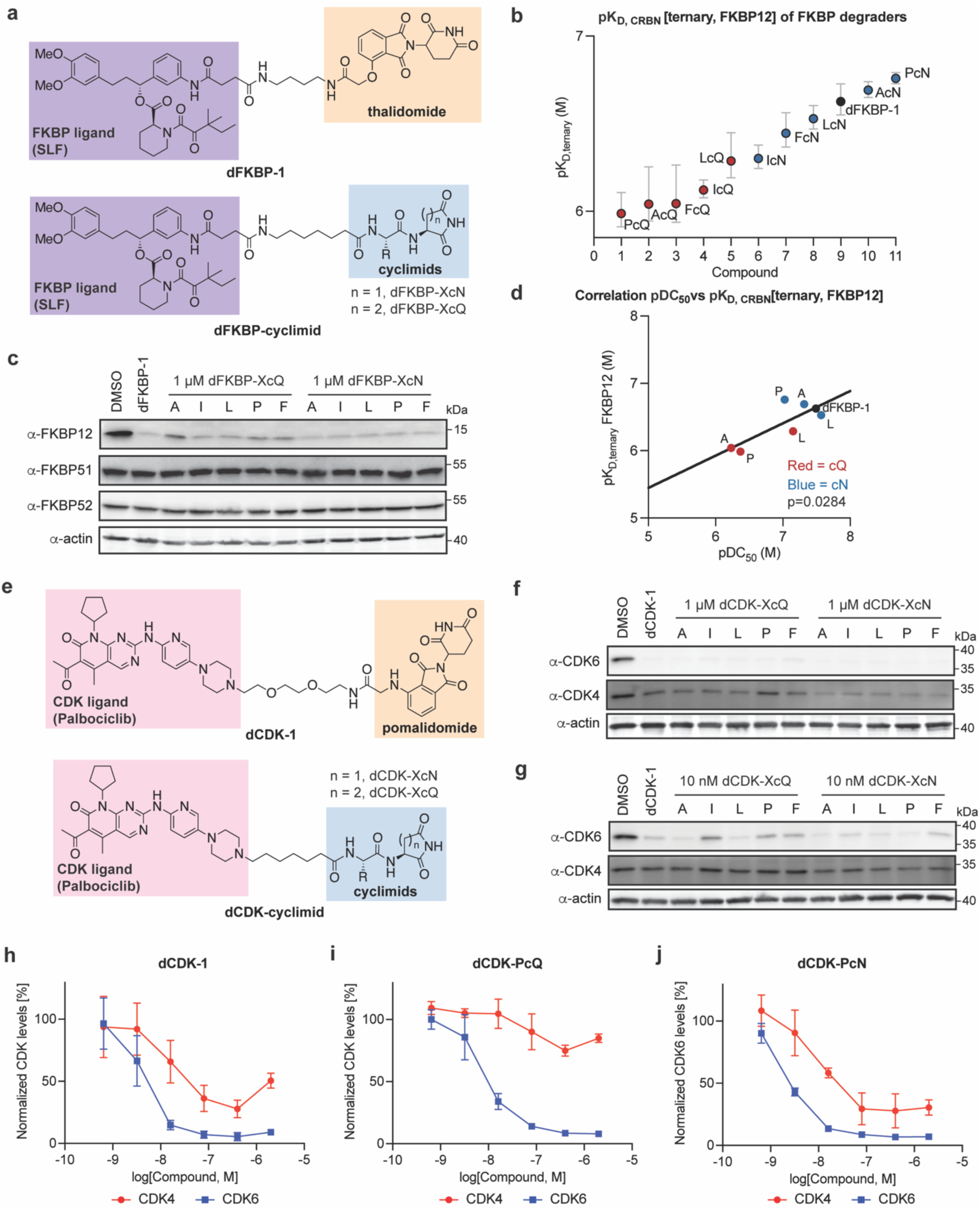
The cyclimid library is readily applicable to target other proteins for selective degradation. (**a**) Structure of dFKBP-1 and cyclimid-based FKBP degraders, dFKBP-cyclimid. (**b**) pK_D, CRBN_[ternary] values of FKBP degraders against CRBN/DDB1 in the presence of a near-saturating concentration of FKBP12 (1 µM). Data are presented as mean ± SD (n = 2 technical replicates). (**c**) Western blot of FKBP12, FKBP51, and FKBP52 levels after treatment of HEK293T cells for 24 h with 10 μM dFKBP-1 or each dFKBP-cyclimid. (**d**) Correlation between pDC_50_ values of FKBP12 degradation and biochemical pK_D, CRBN_[ternary] values. pDC_50_ values of FKBP12 degradation were obtained by quantification of FKBP12 levels relative to DMSO control. Corresponding blot images and quantifications are shown in Extended Data Fig. 8 and uncropped blot images are supplied in Supplementary Fig. 4. (**e**) Structure of dCDK-1 and cyclimid-based CDK degraders, dCDK-cyclimid. (**f**) Western blot of CDK4 and CDK6 levels after treatment of Jurkat cells for 4 h with 1 μM dCDK-1 or each dCDK-cyclimid. (**g**) Western blot of CDK4 and CDK6 levels after treatment of Jurkat cells for 4 h with 10 nM dCDK-1 or each dCDK-cyclimid. (**h–j**) Degradation of CDK4 (red) and CDK6 (blue) after treatment of Jurkat cells for 4 h with (**h**) dCDK-1, (**i**) dCDK-PcQ, or (**j**) dCDK-PcN over a dose-response range of 0.64 nM to 2 μM. Quantification of CDK4 and CDK6 levels were calculated relative to DMSO control. Data in (**h**)–(**j**) are presented as mean ± SD (n = 3 biologically independent samples). Corresponding blot images and quantifications are shown in Extended Data Fig. 9 and uncropped blot images are supplied in Supplementary Fig. 5. All western blot data are representative of at least three independent replicates. For uncropped western blot images, see Supplementary Figure 2.

### Ligand-dependent dynamics of ternary complex dissociation and degradation with the cyclimids

The kinetic properties of PROTAC-mediated ternary complexes, such as their dissociative half-life (*t*_1/2_), have also been shown to influence target protein degradation efficiency, with longer *t*_1/2_ values shown to induce more efficient target degradation, at least for BRD4-targeted PROTACs.^26^ We thus measured the off-rates (*k*_off_) of ternary complexes between His_6_-CRBN/DDB1 and each bromodomain of BRD4 for a subset of the cyclimid degraders (Fig. 4a). Incubation of His_6_-CRBN/DDB1 and GST-BRD4(BD1) or BRD4(BD2) (labeled with AF488-anti-His_6_ antibody and CoraFluor-1-labeled GST antibody, respectively) with concentrations of bifunctional degraders equal or near to their respective K_D, CRBN_[ternary] values (see **Supplementary Table 1**) resulted in the highest assay sensitivity. In the first set of experiments, BRD4(BD1) or BRD4(BD2) dissociation from the ternary complex was initiated by adding JQ1, and dissociation kinetics of the ternary complex with JQ1-AcQ/AcN, JQ1-YcQ/YcN, JQ1-ScQ/ScN, JQ1-GcQ/GcN, or dBET6 were monitored using TR-FRET. These four pairs of cyclimid degraders were selected based on their biochemical selectivity for ternary complexes with BD1 versus BD2. For ternary complexes consisting of His_6_-CRBN/DDB1 and BRD4(BD1), we observed dissociation half-lives (*t*_1/2_) varying from ≤ 30 sec (the time from sample injection to the first measurement) to 270 sec across the degraders tested (Fig. 4b). JQ1-YcQ and JQ1-YcN yielded the most stable and long-lived complexes, consistent with their high positive cooperativity for BRD4(BD1) (Fig. 4c, **Supplementary Table 2**). With BRD4(BD2), however, all ternary complexes exhibited fast dissociation kinetics (*t*_1/2_ ≤ 30 sec) regardless of the choice of degrader, which aligns with the negative cooperativity observed for most of the cyclimid degraders with this bromodomain (Fig. 4b, **Supplementary Table 2**). A reversed dissociation mechanism was also investigated, in which His_6_-CRBN/DDB1 dissociation from the degrader-mediated ternary complex with each bromodomain of BRD4 was triggered by the addition of lenalidomide (Extended Data Fig. 6a). Similarly, ternary complexes with BRD4(BD1) displayed much slower dissociation kinetics relative to ternary complexes with BRD4(BD2) (Extended Data Fig. 6b–c).

**Fig. 6.**
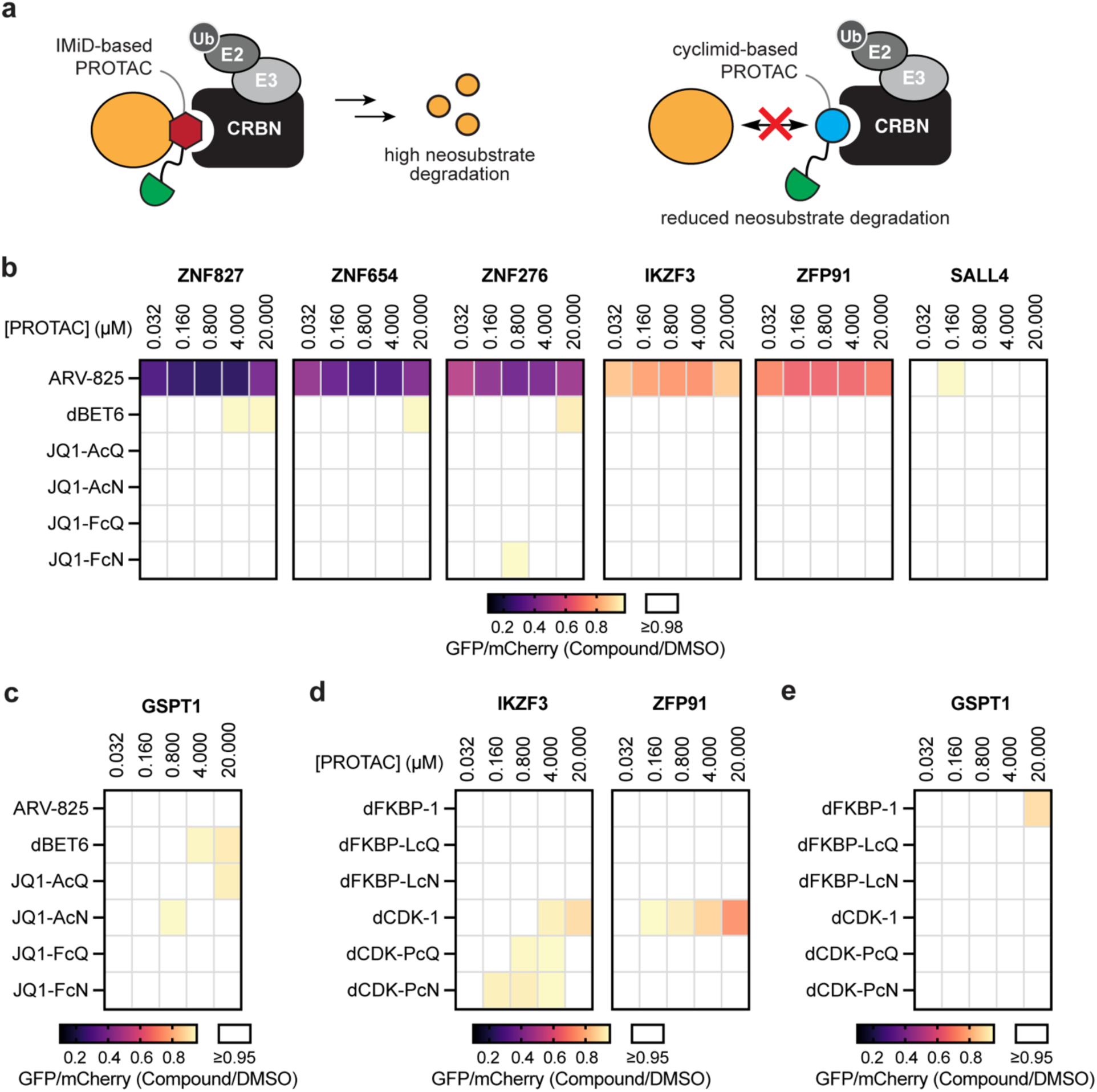
Cyclimid degraders do not recruit known IMiD neosubstrates. (**a**) Schematic of the proposed reduced off-target degradation by the cyclimid-based degraders, in contrast to the IMiD-based degraders. (**b**) Degradation of pomalidomide-sensitive zinc-finger (ZF) degrons in cells by the IMiD- and cyclimid-based BRD4 degraders over a dose-response range of 32 nM to 20 μM. U2OS cells stably expressing 6 ZF degrons fused to GFP were treated with PROTACs followed by flow cytometry to assess ZF degradation. (**c**) Degradation of GSPT1 in cells using IMiD- and cyclimid-based BRD4 degraders. HEK293T cells stably expressing GSPT1-GFP were treated with PROTACs followed by flow cytometry to assess GSPT1 degradation. (**d**) Degradation of pomalidomide-sensitive ZF degrons in cells by IMiD- or cyclimid-based FKBP and CDK degraders over a dose-response range of 32 nM to 20 μM. U2OS cells stably expressing 2 ZF degrons fused to GFP were treated with degraders followed by flow cytometry to assess ZF degradation. (**e**) Degradation of GSPT1 in cells using IMiD- and cyclimid-based FKBP and CDK degraders. HEK293T cells stably expressing GSPT1-GFP were treated with PROTACs followed by flow cytometry to assess GSPT1 degradation. Data in (**b**)–(**e**) are presented as mean (n = 3 biologically independent samples).

As a part of our investigation into the dissociation kinetics and ternary complex formation induced by these BRD4 degraders, the time- and concentration-dependent degradation kinetics of each bromodomain of BRD4 were determined for JQ1-YcQ and dBET6 (Fig. 4d–e). JQ1-YcQ achieved its maximum level of degradation (D_max_) faster with BRD4(BD1) than with BRD4(BD2) (1.5 h for BD1; 3 h for BD2), which indicates that the more favorable ternary complex formation with BD1 than BD2, together with the longer off-rate for BD1 versus BD2, are likely driving the observed difference in degradation dynamics. In contrast, the time required to reach its maximum degradation efficiency with dBET6 was comparable for both bromodomains of BRD4, which reflects the slight difference in the cooperativity and the off-rates of dBET6-induced ternary complexes with BRD4(BD1) or BRD4(BD2). These data provide further evidence that the “occupancy” of cyclimid degraders in a ternary complex (*i.e*., how long the target protein remains tethered to CRBN by the degraders) is another useful parameter in evaluating the degradation activity of the cyclimid degraders, together with K_D, CRBN_[ternary] values.

### The cyclimid library is readily applicable to target other proteins for selective degradation

To further demonstrate the generalizability of the cyclimid platform and the robustness of our TR-FRET-based biochemical approach, we expanded the substrate scope of the cyclimid library beyond the model substrate BRD4. We synthesized a set of cyclimid-based degraders for FKBP, a protein family belonging to the immunophilins, by substituting the thalidomide moiety of dFKBP-1^15^ with each of the ten cyclimids (Fig. 5a). The selection of cyclimids installed in FKBP degraders was based on our observations that nonpolar aliphatic and aromatic cyclimids generally demonstrated superior cellular degradation ability and binding affinity against CRBN compared to cyclimids with polar amino acids at the N–1 position. We additionally included proline because of its unique “bent” structure relative to the rest of the cyclimids. SLF,^47^ an analog of immunosuppressant FK506, is a high-affinity ligand for FKBP12, which has been exploited in IMiD-based degraders like dFKBP-1. As with the evaluation of BRD4 degrader-mediated ternary complexes, using SLF-FITC as a tracer, TR-FRET assays were performed to determine K_D, CRBN_[ternary, FKBP12] of dFKBP-cyclimid degraders and dFKBP-1 by measuring their binding towards CoraFluor-1-labeled His_6_-CRBN/DDB1 in the presence of a near-saturating concentration of recombinant FKBP12 (Extended Data Fig. 7). The dFKBP-cyclimid degraders, like the cyclimid-based BRD4 degraders, displayed diverse K_D, CRBN_[ternary] values, which were greatly altered by the choice of amino acid installed at the N–1 position (Fig. 5b, **Supplementary Table 3**).

Due to the highly homologous binding domain of FKBPs, the FKBP-12 binder SLF has been shown to bind to other FKBP isoforms in the cell.^47^ The degradation activity of the dFKBP-cyclimid library was thus assessed against three members of the FKBP family – FKBP12, FKBP51, and FKBP52 – to determine the generality and selectivity of the cyclimid degraders. With the treatment of HEK293T cells at 1 μM, dFKBP-cyclimid degraders were as effective as the IMiD-based dFKBP-1 in promoting degradation of FKBP12, but did not induce degradation of FKBP51 and FKBP52 (Fig. 5c). Rescue of FKBP12 degradation was observed when cells were pre-treated with a proteasome inhibitor, a neddylation inhibitor, or excess lenalidomide to block CRBN engagement (Extended Data Fig. 8a). While the cyclimid library retains its unique ability to form various ternary complex configurations and to access the productive formation for degradation, each cyclimid revealed a differential potency when embedded in dFKBP structure in comparison with its counterpart BRD4 degrader. For instance, the cyclimid AcQ afforded one of the most potent BRD4 degraders (Fig. 3b, **Supplementary Table 1**), whereas LcQ was the most effective at FKBP12 degradation among the cQ series (Fig. 5c, **Supplementary Table 3**). To assess the relationship between degradation potency and the observed K_D, CRBN_[ternary] values for dFKBP-cyclimids, we determined DC_50_ values for a subset of dFKBP degraders (Extended Data Fig. 8b–d, **Supplementary Table 3**) and correlated these with pK_D, CRBN_[ternary] values. A statistically significant correlation was also observed for the dFKBP-cyclimid series, further reinforcing the utility of K_D, CRBN_[ternary] values as a quantitative indicator for predicting the degradation efficacy in cells (p-value=0.028, Fig. 5d).

We additionally constructed another set of cyclimid-based degraders for the Ser/Thr cell cycle kinases CDK4 and CDK6 to evaluate whether the cyclimids can provide a selective degrader by replacement of the pomalidomide moiety of dCDK-1 (Fig. 5e).^48,49^ The same set of the cyclimids as FKBP degraders was selected when constructing a series of CDK degraders. The dCDK-cyclimid library all facilitated complete degradation of CDK6 at 100 nM, with several degraders being functional at 10 nM (Fig. 5f–g, Extended Data Fig. 9a). Interestingly, the hook effect resulting in reduced degradation of CDK6 can be observed with the cQ-cyclimid degraders and the IMiD-based dCDK-1 at 10 μM, but not with the cN-cyclimid degraders, highlighting an advantageous feature of screening both cN- and cQ-cyclimids as warheads in TPD (Extended Data Fig. 9b). The addition of a proteasome inhibitor, a neddylation inhibitor, or excess lenalidomide completely inhibited CDK4 and CDK6 degradation by a cyclimid degrader, dCDK-PcN, or dCDK-1 (Extended Data Fig. 9c). While most dCDK-cyclimids conferred partial CDK4 depletion at 1 μM, dCDK-PcQ showed a suppressed level of CDK4 degradation relative to other CDK degraders (Fig. 5f). A deeper concentration-dependent profiling over a dose-response range of 0.64 nM to 2 μM with dCDK-PcQ, dCDK-PcN, and dCDK-1 revealed that all three degraders exhibited similar DC_50_ values against CDK6 (DC_50_ = 2–9 nM), whereas dCDK-PcN and dCDK-1, but not dCDK-PcQ, had a significant degradation activity against CDK4 (dCDK-1 and dCDK-PcN: DC_50_ = 10–20 nM with D_max_ = 72 %, Fig. 5h**, j**; dCDK-PcQ: DC_50_ = 89 nM with D_max_ = 25 %, Fig. 5i, **Supplementary Table 4**). Therefore, dCDK-PcQ is remarkably selective towards CDK6 in cellular degradation assays. Intriguingly, the cN-cyclimid series of degraders tend to show superior degradation efficacy to the cQ-cyclimid series, regardless of the choice of a target protein (**Supplementary Table 1, 3–4**). These data indicate that the structurally flexible and versatile cyclimid library can readily tune target selectivity and offers an orthogonal, tunable, and efficient platform for optimizing target degradation and selectivity of heterobifunctional degraders through induction of distinct ternary complexes.

### Cyclimid degraders do not recruit known IMiD neosubstrates

One limitation pertaining to the use of IMiD-based scaffolds in bifunctional degraders for TPD is their potential for off-target toxicity, due to the inherent neosubstrate recruitment by the IMiD warhead, including the teratogenic factor SALL4^50,51^ or the essential translation termination factor GSPT1.^13^ As the cyclimids and the IMiDs promote differentiable binding profiles with CRBN, we expected to observe differentiation in their corresponding neosubstrate recruitment profiles (Fig. 6a). We thus evaluated whether the cyclimid-based bifunctional degraders mitigate the off-target degradation of known IMiD-dependent neosubstrates, adapting a previously reported target profiling platform.^52,53^

Specifically, to assess the off-target properties of the cyclimids, U2OS cells stably expressing several zinc-finger (ZF) degrons fused to GFP^52^ were treated in dose-response with the selected cyclimid degraders, together with the IMiD-based degraders dBET6 and ARV-825,^15,16^ and the degradation of each ZF degron was monitored by flow cytometry (Fig. 6b, Extended Data Fig. 10a). Evaluation of known C2H2 zinc finger targets of the IMiDs, including IKZF3 and ZFP91,^10,11,54^ revealed their fast and rapid degradation by IMiD-based degrader ARV-825. By contrast, almost no degradation of the same off-targets was observed with the cyclimid degraders (D_max_ ≤ 3%), confirming their high capability of reducing unwanted off-target interactions. A second IMiD-based degrader dBET6 also exhibited minimal off-target degradation comparable to the cyclimid degraders (D_max_ ≈ 6%). The off-target properties of the cyclimids were further explored with HEK293T cells stably expressing GSPT1-GFP. After treating the cell line with the same set of degraders, both the cyclimid and the IMiD-based degraders considerably reduced the undesired degradation of GSPT1 (D_max_ ≤ 5% for the cyclimids; D_max_ ≤ 9% for dBET6; D_max_ ≤ 1% for ARV-825, Fig. 6c, Extended Data Fig. 10b). Some cyclimid degraders from the dFKBP and dCDK degrader library, dFKBP-LcQ/LcN and dCDK-PcQ/PcN, were additionally assessed for promiscuity. Using U2OS cells stably expressing ZF degrons of IKZF3 and ZFP91 fused with GFP, as well as HEK293T cells stably expressing GSPT1-GFP, we tested for off-target degradation of these proteins with the dFKBP- and dCDK-cyclimids (Fig. 6d–e). The cyclimid degraders again did not affect the level of those IMiD neosubstrates (D_max_ ≤ 1% for dFKBP-LcQ; D_max_ ≤ 4% for dFKBP-LcN; D_max_ ≤ 6% for dCDK-PcQ; D_max_ ≤ 8% for dCDK-PcN), demonstrating that the cyclimid ligands are unlikely to promote degradation of IMiD neosubstrates, thereby reducing unsought off-target degradation. Further support for the high target selectivity in depleting CDK6 from the cyclimid degraders dCDK-PcQ and dCDK-PcN was provided by unbiased global proteomics (Extended Data Fig. 10c–e).

Encouraged by the dramatically reduced off-target effects of the cyclimids, we characterized their pharmacokinetic properties to inform their translatability to *in vivo* systems. The major limitations of the IMiDs are their proneness to rapid epimerization of the stereocenter in the glutarimide and their hydrolytic instabilities.^55^ By contrast, the cyclimids offer advantages in reducing both destruction pathways. Several monofunctional and bifunctional cyclimid compounds were evaluated for metabolic stability against liver S9 fractions and *in vitro* permeability in a Caco-2 transwell assay, and compared to thalidomide, lenalidomide, or dBET6 (Extended Data Fig. 10f–g). The cyclimid-based degraders were found to have similar metabolic stability and *in vitro* cell permeability characteristics as the IMiD-based degrader dBET6. Monofunctional, JQ1-independent cyclimids showed superior and more favorable metabolic stability than IMiDs. Thus, the cyclimids represent a unique class of CRBN warheads for use in TPD, which possess minimal IMiD-sensitive off-target degradation and promising biochemical and cellular properties to accelerate degrader development for mechanistic studies and translational discovery programs.

## Discussion

Here we report that the cyclimids, a new class of CRBN binders inspired by the C-terminal cyclic imide degron of CRBN, possess distinct modes of interaction with CRBN, thereby enabling efficient sampling of ternary complex space when employed as CRBN ligands in heterobifunctional degraders. A rich diversity of ternary complex structures induced by the cyclimids, combined with their easily translatable properties to target different proteins, offer a facile approach for screening target degradation and selectivity in bifunctional degraders. The structural versatility of the cyclimids enables rapid and efficient screening and optimization to maximize the formation of productive ternary complexes^22,56^ tailored for individual target proteins. An in-depth investigation of cyclimid degraders in binary and ternary complexes across two different substrates (BRD4 and FKBP12) using CoraFluor TR-FRET technology revealed that K_D, CRBN_[ternary] values can serve as a quantitative indicator to predict degradation potency of the cyclimid degraders.

Our work establishes a novel concept for TPD in the ability to rapidly and systematically screen a series of CRBN warheads based on the C-terminal cyclic imide degron. This concept is readily enabled by the cyclimids and represents a complementary approach to the concept of “linkerology” that is widely utilized to discover effective degraders.^36,57,58^ While we report here the use of the 20 canonical amino acids, a natural future extension is to noncanonical amino acids (similar to the strategies employed for developing second-generation VHL-ligands^59^), which will further expand the scope and impact of a warhead screening approach. Cumulatively, the systematic screening of the structure–activity relationship of the CRBN warhead, as we demonstrated with the comprehensive evaluation of over 60 bifunctional degraders, opens a new concept in screening the E3 ligase-binding warhead during degrader development.

The cyclimid library represents a unique class of CRBN-binding ligands for TPD applications, further expanding the current toolbox of emerging potential substitutes^60,61^ for the IMiDs. Degradation of IMiD-sensitive off-targets^51,62^ was not observed with the cyclimid degraders evaluated here, providing evidence that the cyclimids may permit the design of clinically favorable CRBN-targeting therapeutics that avoid long-term side effects for patients. The peptidic properties of the cyclimids may be further harnessed by exploiting the ability of peptide-based molecules to interact with endogenous proteins, such as enzymes and receptors, that recognize specific amino acid sequences or post-translational modifications. These therapeutic strategies may realize desirable drug properties such as tissue-specific targeting and delivery. In light of the diverse suite of ternary complex structures accessible with the cyclimid-based bifunctional degraders, exploring non-peptidic structures will further diversify the cyclimid library and enhance desirable drug-like properties. CRBN-degron-inspired monofunctional degraders may also potentially emerge from these efforts in the future.

We additionally provide the first and most comprehensive evidence that the cyclimids preferentially engage CRBN as the 5-membered aspartimide ligand, which is inactive when installed as the thalidomide derivative. These properties from the cyclimids are in stark contrast to the properties observed with the corresponding thalidomide derivatives. Cyclization of asparagine is more prevalently observed than cyclization of glutamine under physiological conditions due to the formation of the 5-membered aspartimide being kinetically more favorable than the 6-membered glutarimide.^64^ This observation implies that cyclic aspartimides may represent a more dominant degron motif for the thalidomide-binding domain of CRBN than cyclic glutarimides, and CRBN may be more receptive to 5-membered cyclic aspartimides. Focusing on functional differences between the two classes of the cyclimids bearing C-terminal cN or cQ, the cyclimid degraders with 5-membered cN generally exhibited higher affinities to CRBN than their 6-membered cQ counterparts. While one study found that cN and cQ-based peptides exhibited similar engagement with the human thalidomide-binding domain,^63^ the discrepant observation may be attributed to differences in the binding affinity measured by engagement of the thalidomide-bindong domain alone versus the full-length CRBN/DDB1 employed herein. Regardless of the target protein chosen (*e.g*., BRD4, FKBP12, and CDK4/6), the cN-cyclimid degraders largely showed superior degradation efficacy than the cQ-cyclimid series. The higher engagement to CRBN of the cN-cyclimids and the better degradation efficacy when embedded in a heterobifunctional degrader may stem from the more biologically relevant cyclic aspartimide structure. To further illuminate the biological relevance of the cyclimids with CRBN biology, the mechanism of formation of C-terminal cyclic imides – when, where, and how these modifications are formed in the proteome – will need to be elucidated. With the maturation of studies mapping C-terminal cyclic imide sites and substrates across the proteome, the cyclimids may also provide an exciting avenue for designing small molecules that can further tap into native CRBN functions within the proteome.

## Extended Data and Supplementary Information are available for this paper

### Acknowledgments

We thank Y. Amako, D. Miyamoto, and V. Dippon for helpful discussions, and S. Trager and Z. Niziolek from the Harvard University Bauer Core for technical support. His_6_-CRBN/DDB1 was a generous gift from Bristol Myers Squibb. Support from the Ono Pharma Foundation (C.M.W.), Camille–Dreyfus Foundation (C.M.W.), Blavatnik Biomedical Accelerator Grant (C.M.W.), and Japan Society for the Promotion of Science (S.I.), and National Science Foundation graduate research fellowship (N.C.P., H.A.F.) are gratefully acknowledged.

### Author contributions

S.I., C.-F.C., N.C.P., N.V., and S.F. designed and synthesized compounds. S.I. and N.C.P. designed and performed the evaluation of the binary and ternary complex engagement with compounds by TR-FRET assay and associated assay validation. S.I., N.C.P., W.X., and S.F. designed and performed degradation assay. W.X. generated the GSPT1 reporter cell line, designed and performed off-target evaluation experiments with degraders. H.A.F. prepared His_6_-FKBP12 used in TR-FRET assays. C.M.W., R.M., and S.I. conceived of the project. S.I. drafted the manuscript, and all authors jointly discussed the results and edited the manuscript.

### Author information

All data are available in the main text or the Supplementary Information. Proteomics data have been deposited to the PRIDE repository with the dataset identifier PXD041494. Source data are provided with this paper. Correspondence should be addressed to C.M.W. and R.M.

### Methods

#### Cell culture, transfection protocols, and collection

Cells were cultured in DMEM or RPMI 1640 supplemented with 10% heat-inactivated fetal bovine serum (FBS) and 1× penicillin-streptomycin. Cells were grown at 37 °C in a humidified atmosphere with 5% CO_2_. Mycoplasma testing was performed regularly for all cell lines to check for contamination. For the collection of cell pellets, cells were dissociated and collected by two PBS washes and centrifugation at 500 × g, 24 °C, 3 min, followed by a final PBS wash of the pellets. The pellets were flash frozen with liquid nitrogen and stored at –80 °C until use.

#### Western blotting procedures

Unless otherwise noted, cells were lysed by probe sonication (5 sec on, 3 sec off, 15 sec in total, 11% amplitude) in 1–2% SDS in 1× PBS and cleared by centrifugation at 21,000 × g, 4 °C, 10 min. As needed, a BCA assay was performed to determine the protein concentration of lysates and the concentration was adjusted using lysis buffer. 5× SDS-PAGE loading buffer (5% (v/v) β-mercaptoethanol, 0.02% (w/v) bromophenol blue, 30% (v/v) glycerol, 10% (w/v) SDS/ 250 mM Tris pH 6.8) was added to the protein samples to a final concentration of 1× and the samples were heated at 95 °C for 5 min. Protein samples (8–15 μL per lane) were loaded on NuPAGE 3–8% Tris-Acetate precast gels or Criterion XT Tris-Acetate precast gels for high molecular weight proteins, 6/12% Tris-Glycine gels or 4–15% Criterion™ TGX™ precast gels for medium molecular weight proteins, or 16.5% Mini-PROTEAN^®^ Tris-Tricine gels or 15% Criterion Tris-HCl protein gels for low molecular weight proteins. Gels were transferred to membranes using the Invitrogen iBlot 2 dry blotting system and iBlot 2 nitrocellulose transfer stacks, using program P0 (1 min at 20 V, 4 min at 23 V, 2 min at 25 V) for most proteins and 8 min at 25 V for high molecular weight proteins. Membranes were stained with Ponceau S solution to visualize transfer and protein loading. After being blocked with 5% milk or BSA in TBST at 24 °C for 1 h, the membranes were incubated with primary antibodies at 24 °C for 1 h or at 4 °C for 1 to 24 h. Membranes were washed (3 × 5 min) with TBST and incubated with secondary antibodies at 24 °C for 1 h. Membranes were washed (3 × 5 min) with TBST and the results were obtained by chemiluminescence and/or IR imaging using Azure 600 or c600.

#### BRD4 degradation assay

1.5×10^6^ HEK293T cells were seeded in DMEM supplemented with 10% FBS and 1× penicillin-streptomycin and incubated at 37 °C, 5% CO_2_ for 30 min, then treated with compounds of interest and incubated at 37 °C, 5% CO_2_ for 4 h prior to collection and lysis. If noted, cells were treated with 1 μM MLN4924 for 1 h before treatment with compounds following the initial 30 min incubation. All compounds were dissolved in DMSO, and the final DMSO concentration after addition of the compound to the cells did not exceed 0.2% v/v.

#### FKBP12, FKBP51, and FKBP52 degradation assay

1.5×10^6^ HEK293T cells were seeded in DMEM supplemented with 10% FBS and 1× penicillin-streptomycin and incubated at 37 °C, 5% CO_2_ for 30 min, then treated with compounds of interest and incubated at 37 °C, 5% CO_2_ for 18 h prior to collection and lysis. If noted, cells were treated with 1 μM MLN4924 for 1 h before treatment with compounds following the initial 30 min incubation. All compounds were dissolved in DMSO and the final DMSO concentration after addition of the compound to the cells did not exceed did not exceed 0.2% v/v.

#### CDK4 and CDK6 degradation assay

1.5×10^6^ Jurkat cells were seeded in RPMI supplemented with 10% FBS and 1× penicillin-streptomycin and incubated at 37 °C, 5% CO_2_ for 30 min, then treated with compounds of interest and incubated at 37 °C, 5% CO_2_ for 24 h prior to collection and lysis. If noted, cells were treated with 1 μM MLN4924 for 1 h before treatment with compounds following the initial 30 min incubation. All compounds were dissolved in DMSO and the final DMSO concentration after addition of the compound to the cells did not exceed did not exceed 0.2% (v/v).

#### Cloning, overexpression, and purification of His_6_-FKBP12

The cloning, overexpression, and purification of His_6_-FKBP12 were adapted from Flaxman and co-workers.^65^ In brief, Rosetta 2 (DE3) chemically competent cells transformed with the pGEX2T-His_6_-FKBP12 plasmid were used to inoculate a culture of autoclaved Luria Bertani broth with 100 μg/mL ampicillin (50 mL). The culture was grown at 30 °C for 15–18 h with shaking at 200 rpm. The saturated culture was then diluted 1:100 into autoclaved Luria Bertani Broth (750 mL) with 100 μg/mL ampicillin in a 2 L baffled flask. The OD600 of the culture was monitored until the OD600 measured between 0.3–0.6, at which point protein expression was induced with 0.3 mM IPTG. After the addition of IPTG, the temperature of the incubator was reduced to 25 °C. The culture was incubated with shaking at 200 rpm at 25 °C for 5 h. Cells were harvested by centrifugation at 3,750 rpm (2,249 × g, 24 °C, 15 min). The pellet was re-suspended in lysis buffer (5 mL, 1 mM DTT, 0.5 mg/mL lysozyme/PBS), added protease inhibitor, and lysed by sonication (30 sec on, 10 sec off, 15% amplitude, 5 min total). DNAse (0.8 mg/mL) and RNAse (0.008 mg/mL) were added to the lysate and incubated for 30 min. After the clarification by centrifugation (10,000 x g, 4 °C, 10 min), followed by filtration through a 0.45 μm syringe filter, the clarified lysate was applied to the column with glutathione sepharose resin (1 mL) and shaken at 4 °C for 1–2 h. The unbound lysate was discarded, and the resin was sequentially washed with 1 mM DTT/PBS (20 mL) and thrombin cleavage buffer (5 mL, 0.3 mM CaCl2, 1 mM DTT /20 mM Tris, 100 mM NaCl, pH 6.8). The column was then treated with thrombin (100 units) in thrombin cleavage buffer (5 mL) and incubated with shaking at 4 °C for 15–18 h. The cleaved protein was collected from the column, which was further washed with 1 mM DTT/PBS (20 mL). The remaining GST was eluted with 10 mM reduced glutathione/50 mM Tris, pH 8.0 (5 mL). The thrombin cleavage and following washes were concentrated to 500 μL using a 3 kDa MWCO spin concentrator, with thrombin removal in conjunction with final buffer exchange. The final purification and buffer exchange were performed by SEC using a Superdex 75 GL 10/300 column, run with PBS at a flow rate of 0.5 mL/min. The desired protein was then collected from FPLC fractions and concentrated using a 3 kDa MWCO spin concentrator until the protein concentration was >1 mg/mL. Protein solution was aliquoted, flash frozen with liquid nitrogen, and stored at –80 °C until use.

#### TR-FRET measurements

Unless otherwise noted, experiments were performed in white, 384-well microtiter plates (Corning 3572 or Greiner 781207) in 30 μL assay volume. TR-FRET measurements were acquired on a Tecan SPARK plate reader with SPARKCONTROL software version V2.1 (Tecan Group Ltd.), with the following settings: 340/50 nm excitation, 490/10 nm (Tb), and 520/10 nm (FITC, AF488, GFP) emission, 100 μs delay, 400 μs integration. The 490/10 nm and 520/10 nm emission channels were acquired with a 50% mirror and a dichroic 510 mirror, respectively, using independently optimized detector gain settings unless specified otherwise. The TR-FRET ratio was taken as the 520/490 nm intensity ratio on a per-well basis.

#### Protein labeling for TR-FRET measurements

Anti-His_6_ antibody (Abcam 18184), anti-GST antibody (Abcam 19256), anti-GST V_H_H (ChromoTek ST-250), anti-GFP V_H_H (ChromoTek GT-250), anti-RFP V_H_H (ChromoTek RT-250), anti-rabbit Nano-Secondary (ChromoTek shurbGNHS-1), and His_6_-CRBN/DDB1 complex were labeled with CoraFluor-1-Pfp ester or AF488-Tfp ester (ThermoFisher A37570) as previously described.^33,66^ The following extinction coefficients were used to calculate protein concentration and degree-of-labeling (DOL): IgG *E*_280_ = 210,000 M^−1^cm^−1^, anti-GST V_H_H *E*_280_ = 28,545 M^−1^cm^−1^, anti-GFP V_H_H *E*_280_ = 27,055 M^−1^cm^−1^, anti-RFP V_H_H *E*_280_ = 30,035 M^−1^cm^−1^, anti-rabbit Nano-Secondary *E*_280_ = 24,075 M^−1^cm^−1^, His_6_-CRBN-DDB1 *E*_280_ = 167,000 M^−1^cm^−1^, CoraFluor-1-Pfp *E*_340_ = 22,000 M^−1^cm^−1^, AF488-Tfp *E*_495_ = 71,000 M^−1^cm^−1^. Protein conjugates were snap-frozen in liquid nitrogen, and stored at –80 °C.

#### Determination of equilibrium dissociation constants (*K*_D_ or *K*_D,app_) for fluorescent tracers by TR-FRET

The following conditions have been employed: (i) 1 nM CoraFluor-1-labeled anti-His_6_ antibody, 2.5 nM His_6_-CRBN/DDB1 complex in assay buffer (25 mM HEPES, 150 mM NaCl, 0.5 mg/mL BSA, 0.005% TWEEN-20, pH 7.5), (ii) 2.5 nM CoraFluor-1-labeled His_6_-CRBN/DDB1 complex in assay buffer, (iii) 0.5 nM GST-BRD4(BD1) or GST-BRD4(BD2), 4 nM CoraFluor-1-labeled anti-GST V_H_H in assay buffer, (iv) 2.5 nM CoraFluor-1-labeled anti-His_6_ antibody, 5 nM His_6_-FKBP12 in assay buffer, (v) 1 nM CoraFluor-1-labeled anti-His_6_ antibody, 0.5 nM His_6_-GST-CDK4/GST-CD3 in assay buffer, (vi) 1 nM CoraFluor-1-labeled anti-6xHis antibody, 2 nM His_6_-CDK6/GST-CD1 in assay buffer.

Fluorescent tracers (Thalidomide-FITC for CRBN-DDB1 assays, JQ1-FITC for BRD4 assays, SLF-FITC for FKBP12 assays, Palbociclib-FITC for CDK4/6 assays) were added in serial dilution using a HP D300 digital dispenser (Hewlett-Packard) and allowed to equilibrate for 1 h at room temperature before TR-FRET measurements were taken. For all assays using CoraFluor-1-labeled anti-epitope tag reagents (anti-6xHis antibody, anti-GST V_H_H), nonspecific signal was determined from wells containing the CoraFluor-1-labeled conjugate alone in the absence of the target protein. For assays using CoraFluor-1-labeled His_6_-CRBN/DDB1 complex, nonspecific signal was determined from wells containing 100 µM Lenalidomide-5′-NH_2_. Data were fitted to a One Site – Specific Binding model in Prism 9.

For conditions (i), (iii), and (iv), the *K*_D_ determined from the one-site model was then used in Equation 1 to adjust for a two-site model due to the presence of dimeric GST protein or bivalent anti-6xHis antibody:

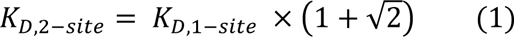

For conditions (v) and (vi), we did not attempt to determine the absolute *K*_D_ values for the Palbociclib-FITC tracer due to the potential presence of higher-order oligomeric complexes. For CDK4/6 experiments, *K*_D,app_ values for Palbociclib-FITC and test compounds are reported.

In subsequent TR-FRET ligand displacement assays, inhibitor *K*_D_ and *K*_D,app_ values have been calculated using Cheng-Prusoff principles, outlined in Equation 2 below:

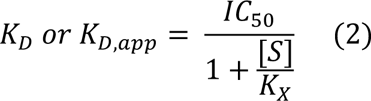

Where IC_50_ is the measured IC_50_ value, [S] is the concentration of fluorescent tracer, and *K*_X_ is the *K*_D_ or *K*_D,app_ of the fluorescent tracer for a given condition [4202581].

#### TR-FRET ligand displacement assays

The following conditions have been employed: (i) 1 nM CoraFluor-1-labeled anti-His_6_ antibody, 2.5 nM His_6_-CRBN/DDB1 complex, and 50 nM Thalidomide-FITC in assay buffer, (ii) 2.5 nM CoraFluor-1-labeled His_6_-CRBN/DDB1 complex, 200 nM Thalidomide-FITC in assay buffer, (iii) 4 nM GST-BRD4(BD1) or GST-BRD4(BD2), 4 nM CoraFluor-1-labeled anti-GST V_H_H, and 20 nM JQ1-FITC in assay buffer, (iv) 2.5 nM CoraFluor-1-labeled anti-His_6_ antibody, 5 nM His_6_-FKBP12, and 60 nM SLF-FITC in assay buffer, (v) 1 nM CoraFluor-1-labeled anti-His_6_ antibody, 2 nM His_6_-GST-CDK4/GST-CD3, and 30 nM Palbociclib-FITC in assay buffer, (vi) 1 nM CoraFluor-1-labeled anti-His_6_ antibody, 2 nM His_6_-CDK6/GST-CD1, and 200 nM Palbociclib-FITC in assay buffer.

In all cases, test compounds were added in serial dilution (1:2 titration, 15-point, c_max_ = 10 μM) using a HP D300 digital dispenser and allowed to equilibrate for 1 h at room temperature before TR-FRET measurements were taken. For conditions (i-iii), the assay floor was determined from wells treated with 10 μM dBET6. For condition (iv), the assay floor was determined from wells treated with 10 μM dFKBP-1. For conditions (v-vi), the assay floor was determined from wells treated with 10 μM Palbociclib. The assay ceiling (top) was defined via a no-inhibitor control. Data were background-corrected, normalized and fitted to a four-parameter dose-response model [log(inhibitor vs. response – Variable slope (four parameters)] in Prism 9, with constraints of Top = 1 and Bottom = 0.

#### Determination of the cooperativity constant (α) for PROTAC-mediated ternary complexes of CRBN/DDB1 with recombinant BRD4 bromodomains and FKBP12

Assays were performed in low-volume, white 384-well plates (ProxiPlate-384 Plus, PerkinElmer) in 10 μL assay volume. The following assay conditions have been employed: (i) 1 nM CoraFluor-1-labeled anti-His_6_ antibody, 2.5 nM His_6_-CRBN/DDB1 complex, 50 nM Thalidomide-FITC, and 1 μM GST-BRD4(BD1) in assay buffer, (ii) 1 nM CoraFluor-1-labeled anti-His_6_ antibody, 2.5 nM His_6_-CRBN/DDB1 complex, 50 nM Thalidomide-FITC, and 1 μM GST-BRD4(BD2) in assay buffer, (iii) 2.5 nM CoraFluor-1-labeled His_6_-CRBN/DDB1 complex, 200 nM Thalidomide-FITC, and 1 μM His_6_-FKBP12 in assay buffer.

In all cases, test compounds were added in serial dilution (1:4 titration, 7-point, c_max_ = 1 μM) using a HP D300 digital dispenser and allowed to equilibrate for 1 h at room temperature before TR-FRET measurements were taken. The assay floor was determined from wells treated with 10 μM dBET6, and the assay ceiling (top) was defined via a no-inhibitor control. Data were background-corrected, normalized and fitted to a four-parameter dose-response model [log(inhibitor vs. response – Variable slope (four parameters)] in Prism 9, with constraints of Top = 1 and Bottom = 0.

The cooperativity constant (α) was calculated via Equation 3 below:

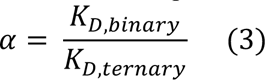

Where *K*_D,binary_ and *K*_D,ternary_ are the respective inhibitor *K*_D_ values in the absence and presence of 1 µM recombinant target protein (e.g. GST-BRD4(BD1), GST-BRD4(BD2), or His_6_-FKBP12).

#### TR-FRET assay to quantify endogenous BRD4 abundance in MDA-MB-231 cell extract after degrader treatment

Assays were run as previously reported.^38^ MDA-MB-231 cells were seeded into 96-well plates (Corning 3903 or Corning 3904) at a density of 20,000 cells/well in 100 μL cell culture medium and allowed to adhere overnight. A HP D300 digital dispenser was used to dispense serial dilutions of test compounds (1:2 titration, 12-point, c_max_ = 1 μM) normalized to 0.1% DMSO. Cells were incubated for 5 h at 37°C and 5% CO_2_ then media was replaced with pre-warmed cell culture medium (150 μL/well) and residual test compound was washed out for 1 h at 37°C and 5% CO_2_. After, media was aspirated and cells were washed with PBS (200 μL/well), followed by the addition of ice-cold lysis buffer (60 μL/well; 25 mM HEPES, 150 mM NaCl, 0.2% v/v Triton X-100, 0.02% TWEEN-20, pH 7.5 supplemented with 1 mM AEBSF hydrochloride (Combi-Blocks SS-7834), 250 U Benzonase (Sigma E1014), and 2 mM DTT). The plate was shaken at 1,000 rpm on an orbital shaker (Boekel Scientific Jitterbug, model 130000) for 10 min. After, 10 μL of 7x control detection mix (7 nM CoraFluor-1-labeled anti-rabbit Nano-Secondary, 700 nM JQ1-FITC final concentrations, prepared in dilution buffer; 25 mM HEPES, 150 mM NaCl, 0.005% TWEEN-20, pH 7.5) was added to four of the eight DMSO-treated control wells, and 10 μL of complete detection mix (3.5 nM rabbit anti-BRD4 antibody, 7 nM CoraFluor-1-labeled anti-rabbit Nano-Secondary, 700 nM JQ1-FITC final concentrations, prepared in dilution buffer) was added to all other wells and allowed to equilibrate for 1 h. The plate was centrifuged at 2,000 × *g* for 1 min then lysate was transferred to a 384-well plate (30 μL × 2 TR-FRET replicates) using an adjustable electronic multichannel pipette (Matrix Equalizer, ThermoFisher 2231), and TR-FRET measurements were taken.

TR-FRET ratios were background-subtracted from the four (eight TR-FRET replicates) DMSO-treated wells containing control detection mix. The average TR-FRET intensity was then normalized to the four (eight TR-FRET replicates) DMSO-treated wells containing complete detection mix for each biological replicate, then data were fitted to a four-parameter dose-response model [log(inhibitor vs. response – Variable slope (four parameters)] in Prism 9, with constraints of Top = 1.

#### TR-FRET assay to quantify BRD4(BD1)-GFP and BRD4(BD2)-GFP in lysates after degrader treatment

2.0×10^5^ HEK293T cells stably expressing either BRD4(BD1)-GFP or BRD4(BD2)-GFP cells^22^ were seeded in DMEM supplemented with 10% FBS and 1× penicillin-streptomycin and incubated at 37 °C, 5% CO_2_ for 30 min, then treated with compounds of interest and incubated at 37 °C, 5% CO_2_ for the indicated time prior to collection and lysis. All compounds were dissolved in DMSO, and the final DMSO concentration after addition of the compound to the cells did not exceed 0.2% v/v. After compound treatment, cells were collected, washed with PBS (500 μL), flash-frozen, and stored at –80 °C until further use.

Frozen cell pellets (∼200,000 cells per pellet) were thawed on ice and to each pellet was added 200 uL ice-cold complete lysis buffer. The cell pellets were briefly vortexed and lysis was allowed to proceed at room temperature for 10 min. After, lysates were transferred to a 384-well plate (30 μL × 2 TR-FRET replicates), followed by the addition of 5 μL of 7x detection mix (28 nM CoraFluor-1-labeled anti-GFP V_H_H in dilution buffer). The plate was allowed to equilibrate for 1 h at room temperature before TR-FRET measurements were taken.

TR-FRET ratios were background-subtracted from wells containing 0.5 mg/mL BSA + detection mix. TR-FRET ratios were then normalized to the DMSO-treated wells for each biological replicate. Data were fitted to a four-parameter dose-response model [log(inhibitor vs. response – Variable slope (four parameters)] in Prism 9, with constraints of Top = 1.

#### Off-rate measurements for degrader-mediated ternary complexes between His6-CRBN/DDB1 and individual GST-tagged bromodomains

Assays were performed in low-volume, white 384-well plates (ProxiPlate-384 Plus, PerkinElmer) in 10 μL assay volume. A master solution of His_6_-CRBN/DDB1 (10 nM), AF488-labeled anti-His_6_ antibody (10 nM; Ab18184), CoraFluor-1-labeled anti-GST antibody (5 nM; Ab19256), and GST-BRD4(BD1) or GST-BRD4(BD2) (10 nM) was prepared and divided into 59 uL aliquots. After, 1 µL of 60x compound stock (in DMSO) was added to achieve the following concentrations: for GST-BRD4(BD1), 10 nM dBET6, 10 nM JQ1-YcQ, 10 nM JQ1-YcN, 20 nM JQ1-AcQ, 10 nM JQ1-AcN, 50 nM JQ1-ScQ, 10 nM JQ1-ScN, 120 nM JQ1-GcQ, 20 nM JQ1-GcN; for GST-BRD4(BD2), 20 nM dBET6, 275 nM JQ1-YcQ, 150 nM JQ1-YcN, 75 nM JQ1-AcQ, 20 nM JQ1-AcN, 150 nM JQ1-ScQ, 25 nM JQ1-ScN, 540 nM JQ1-GcQ, 90 nM JQ1-GcN. These concentrations were chosen explicitly as they are equal to or near the K_D, CRBN_[ternary] values for each degrader (see Supplementary Table 1).

Solutions were allowed to equilibrate for 1 h at room temperature then 3 × 10 uL technical replicates were plated into wells of the 384-well plate before initial *t* = 0 TR-FRET measurements were taken. Following the addition of JQ1 (10 µM) or Lenalidomide-5′-NH_2_ (50 µM), the time-dependent change in TR-FRET signal was recorded over 10 min in 30 s intervals using fixed detector gain settings. Data were background-corrected from wells containing no compound, normalized to the *t* = 0 TR-FRET intensity on a per-compound basis, and fitted to a one phase decay model using Prism 9.

#### Generation of off-target degradation reporter cell lines

U2OS cells stably expressing the 23-aa zinc finger region of IKZF3, SALL4, ZFP91, ZNF276, ZNF654, and ZNF827 were kindly provided by the Choudhary Lab.^52^ The procedures of molecular cloning and cell line generation were adapted from Sievers and coworkers.^53^ The Cilantro2-GSPT1 construct was derived from the degradation reporter vector Cilantro2 and pcDNA3.1-GSPT1. The GSPT1 insert was amplified by overhang PCR using Primer 1 and Primer 2 to add the BsmBI restriction sites. The vector and insert were digested by BsmBI, ligated by Quick Ligation Kit, and transformed into NEB Stable cells. The sequence was validated by Sanger sequencing (Quintara Bio).

For virus production, HEK293T cells in a 10 cm plate were transfected by pMD2.G (6 µg), psPAX2 (4 µg) and Cilantro2-GSPT1 (10 µg) using TransIT-Pro reagent (20 µL) following the manufacturer’s guidelines. Lentiviruses were collected 48 h after transfection, spun down (500 × g, 5 min, 24 °C) and filtered with a 0.45 µm PES filter. For transduction, HEK293T cells in 1 mL DMEM+/– containing 10 µg/mL polybrene were seeded into 6-well plates containing the virus (500 µL) and incubated for 48 h and then selected by 2 µg/mL puromycin for 7 days. The GSPT1 reporter cell line was validated by a test degradation assay using the GSPT1 degrader CC-885.^13^

#### Off-target degradation assay by flow cytometry

Cells were seeded in 96-well plates (3–4×10^4^ cells per well) 1 h before compound treatment. DMSO or each compound at each concentration was added into three separate wells (final DMSO concentration: 0.2%) and incubated at 37 °C, 5% CO_2_ for 24 h. The cells were detached by trypsinization in 0.25% trypsin without phenol red (diluted 2× by PBS), followed by resuspension in DMEM +/– without phenol red. All plates were analyzed by FACSymphony (BD Biosciences) on the high throughput sampler.

Signal from at least 2,000 events per well was acquired, and the fluorescence signals of FITC and PE Texas Red were measured. GFP/mCherry ratios were determined by dividing the geometric mean of FITC signal by that of PE Texas Red signal on BD FACSDiva v9.1. For each condition, the arithmetic mean of ratios among triplicates was calculated and then normalized to that of the DMSO ratios to derive the GFP/mCherry (compound/DMSO) value.

#### Computational modeling

Computational models were generated in MOE version 2022.02. The IMiD ligand from the indicated crystal structure was adapted to the indicated dipeptide and the adapted complex was subjected to energy minimization with Amber10:EHT force field followed by preparation with Protonate 3D.

#### Global quantitative proteomics sample preparation

Global proteomics samples were prepared in biological triplicate for each condition. 2.0×10^6^ Jurkat cells were seeded in 12-well plates and incubated at 37 °C, 5% CO_2_ for 1 h. Small molecules of interest were then added, and cells were incubated at 37 °C, 5% CO_2_ for 4 h. Cells were collected according to the described general procedure, lysed by probe sonication (5 sec on, 3 sec off, 15 sec in total, 11% amplitude) in lysis buffer (8 M urea, 50 mM NaCl, 50 mM HEPES, 1x protease/phosphatase inhibitor cocktail, pH 8.2), and cleared by centrifugation (21,000 × g, 4 °C, 10 min). After protein quantification by BCA protein assay, the lysates were diluted to 0.89 mg/mL with the lysis buffer. The diluted lysates (110 μL) were reduced by the addition of dithiothreitol (final concentration: 5 mM) at 24 °C for 30 min then alkylated by addition of iodoacetamide (final concentration: 15 mM) and incubation in the dark at 24 °C for 30 min. The proteins in the samples were precipitated using methanol-chloroform precipitation. In brief, four volumes of chilled methanol, one volume of chilled chloroform, and three volumes of water were added sequentially to the lysates. The mixture was vortexed and centrifugated at 14,000 × g, 5 min, 4 °C, and the supernatant was aspirated. The protein pellet was washed with three volumes of chilled methanol and centrifugated at 14,000 × g, 5 min, 4 °C, and the resulting precipitated protein was air-dried.

The protein was dissolved in 25 µL 4M urea, 50 mM HEPES, pH 7.4, followed by the addition of 75 µL 200 mM HEPPS, pH 8.0. Lys-C (2.0 µg) was added to the mixture, and the digestion was allowed to proceed at 30 °C for 4 h without rotation. The samples were diluted by the addition of 200 mM HEPPS, pH 8.0 (100 µL), and further digested by trypsin (4.0 µg) at 37 °C for 18 h without rotation. Approximately 50 µg of peptides from each digested sample (107 μL) was taken for labeling with TMTpro 16-plex reagent (17 μL) at 24 °C for 1 h. TMT labeling was quenched by incubation with 5% hydroxylamine (7 µL) at 24 °C for 15 min. The TMT-labeled samples were combined and dried by a vacufuge. The combined, dried sample was resuspended in 1800 µL 0.1% trifluoroacetic acid (TFA) in water and divided into 6 portions of 300 µL solution. Each portion was taken to fractionation using a Pierce high pH reversed-phase peptide fractionation kit (6 columns used in total). The peptides were eluted sequentially by 4% acetonitrile/0.1% triethylamine (TEA) through 20% acetonitrile/0.1% TEA in 1% acetonitrile increments (17 fractions), followed by 25%, 30%, and 50% acetonitrile/0.1% TEA. The first fraction (4% acetonitrile/0.1% TEA) was excluded from LC-MS/MS analysis. The other fractions were concentrated to dryness by a vacufuge. The 6 fractions eluted with the same percentage of acetonitrile from each column were combined and resuspended in 60 µL of 0.1% formic acid prior to LC-MS/MS analysis (19 fractions in total).

#### Proteomics mass spectrometry acquisition procedures for protein level quantitation

Desalted and fractionated samples were resuspended in 0.1% formic acid/water (60 μL per sample). The samples (6–8 μL) were loaded onto a C18 trap column (3 cm, 3 μm particle size C18 Dr. Maisch 150 μm I.D) and then separated on an analytical column (50 cm PharmaFluidics, Belgum) at 0.2 µL/min with a Thermo Scientific Ultimate 3000 system connected in line to a Thermo Scientific Orbitrap Fusion Lumos Tribrid. The column oven temperature was maintained at 35 °C. Peptides were eluted using a multi-step gradient at a flow rate of 0.3 µL/min over 90 min (0–10 min, 5% acetonitrile in 0.1% formic acid/water; 10–72 min, 5–32%; 72–80 min, 32–98%; 80–90 min, 98%). The electrospray ionization voltage was set to 2.2 kV and the capillary temperature was set to 275 °C. Dynamic exclusion was enabled with a mass tolerance of 10 ppm and exclusion duration of 90 sec. MS1 scans were performed over 400–2000 m/z at resolution 120,000. HCD fragmentation was performed on the top ten most abundant precursors exhibiting charge states from two to five at a resolving power setting of 60,000 and fragmentation energy of 37% in the Orbitrap. CID fragmentation was applied with 35% collision energy, and resulting fragments were detected using the normal scan rate in the ion trap.

#### Mass spectrometry data analysis for protein level quantitation

Analysis was performed in Thermo Scientific Proteome Discoverer version 2.4.1.15. The raw data were searched against SwissProt human (*Homo sapiens*) protein database (17 February 2022; 42,279 total entries) and contaminant proteins using the Sequest HT algorithm. Searches were performed with the following guidelines: spectra with a signal-to-noise ratio greater than 1.5; mass tolerance of 20 ppm for the precursor ions and 0.02 Da (HCD) and 0.6 Da (CID) for fragment ions; full trypsin digestion; 2 missed cleavages; variable oxidation on methionine residues (+15.995 Da); static carboxyamidomethylation of cysteine residues (+57.021 Da); static TMTpro labeling (+304.207 Da) at lysine residues and N-termini. The TMT reporter ions were quantified using the Reporter Ions Quantifier node and normalized to the total peptide amount. Peptide spectral matches (PSMs) were filtered using a 1% false discovery rate (FDR) using Percolator. PSMs were filtered to PSMs in only one protein group with an isolation interference under 70%. For the obtained proteins, the data were filtered to include master and master candidate proteins with high protein FDR confidence and exclude all contaminant proteins. The ratios and p-values were obtained from Proteome Discoverer (p-values were calculated by one-way ANOVA with TukeyHSD post-hoc test).

---

**Extended Data Fig. 1.**
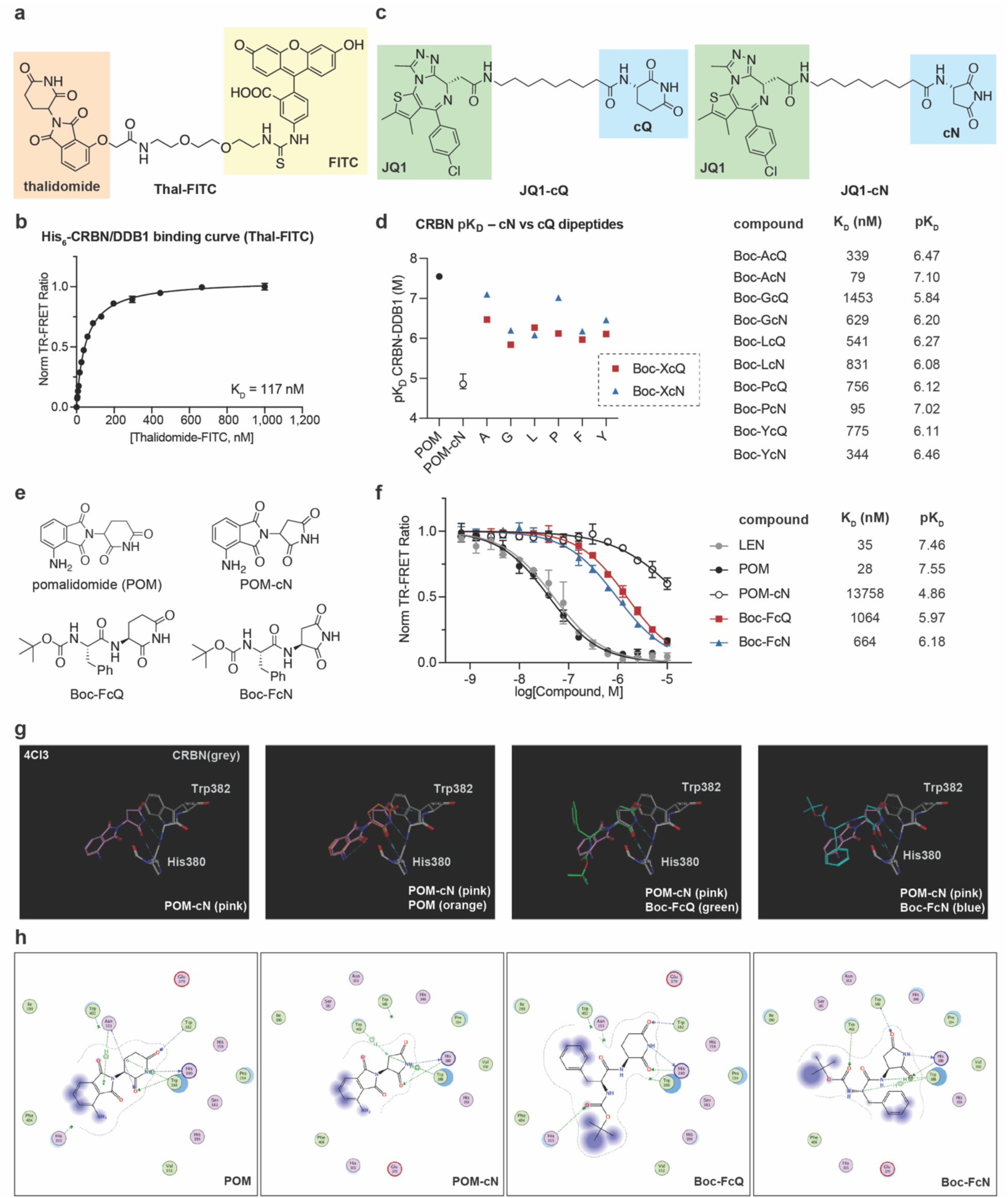
TR-FRET ligand displacement assay platform to quantitatively measure small molecule target engagement with His_6_-CRBN/DDB1, BD1 and BD2 domains of BRD4. (**a**) Structure of Thal-FITC, a tracer used to assess the engagement with CRBN/DDB1. (**b**) Saturation binding of Thal-FITC to His_6_-CRBN/DDB1 labeled with CoraFluor-1-anti-His_6_ conjugates. Data are presented as mean ± SD (n = 3 technical replicates). (**c**) Structures of JQ1-cQ and JQ1-cN. (**d**) Comparison between monofunctional cQ- and cN-cyclimids regarding pK_D_ values (equilibrium dissociation constants; K_D_) against CRBN/DDB1. The cN-cyclimids generally bind to CRBN/DDB1 more tightly than their cQ counterparts. (n = 3 technical replicates, error bars, which fall within symbols, represent mean ± SD). (**e**) Structures of pomalidomide (POM), POM-cN, Boc-FcQ, and Boc-FcN. (**f**) Dose-titration of the indicated monofunctional compounds in TR-FRET ligand displacement assays with His_6_-CRBN/DDB1 and the determined K_D_ values of the indicated compounds against the His_6_-CRBN/DDB1 complex. Data are presented as mean ± SD (n = 3 technical replicates). (**g**) Structural models of POM, POM-cN, Boc-FcQ, and Boc-FcN, built from the crystal structure of the CRBN/DDB1 complex bound to pomalidomide (PDB: 4CI3). Zoom-in of the thalidomide-binding pocket of CRBN with POM-cN (pink), and overlays of POM-cN with POM (orange), Boc-FcQ (green), or Boc-FcN (blue). (**h**) Sketch of POM, POM-cN, Boc-FcQ, and Boc-FcN and their interactions with residues in the thalidomide-binding pocket of CRBN. POM-cN is incapable of forming hydrogen bonds with Trp 382 of CRBN, in contrast to POM, Boc-FcQ, and Boc-FcN.

**Extended Data Fig. 2.**
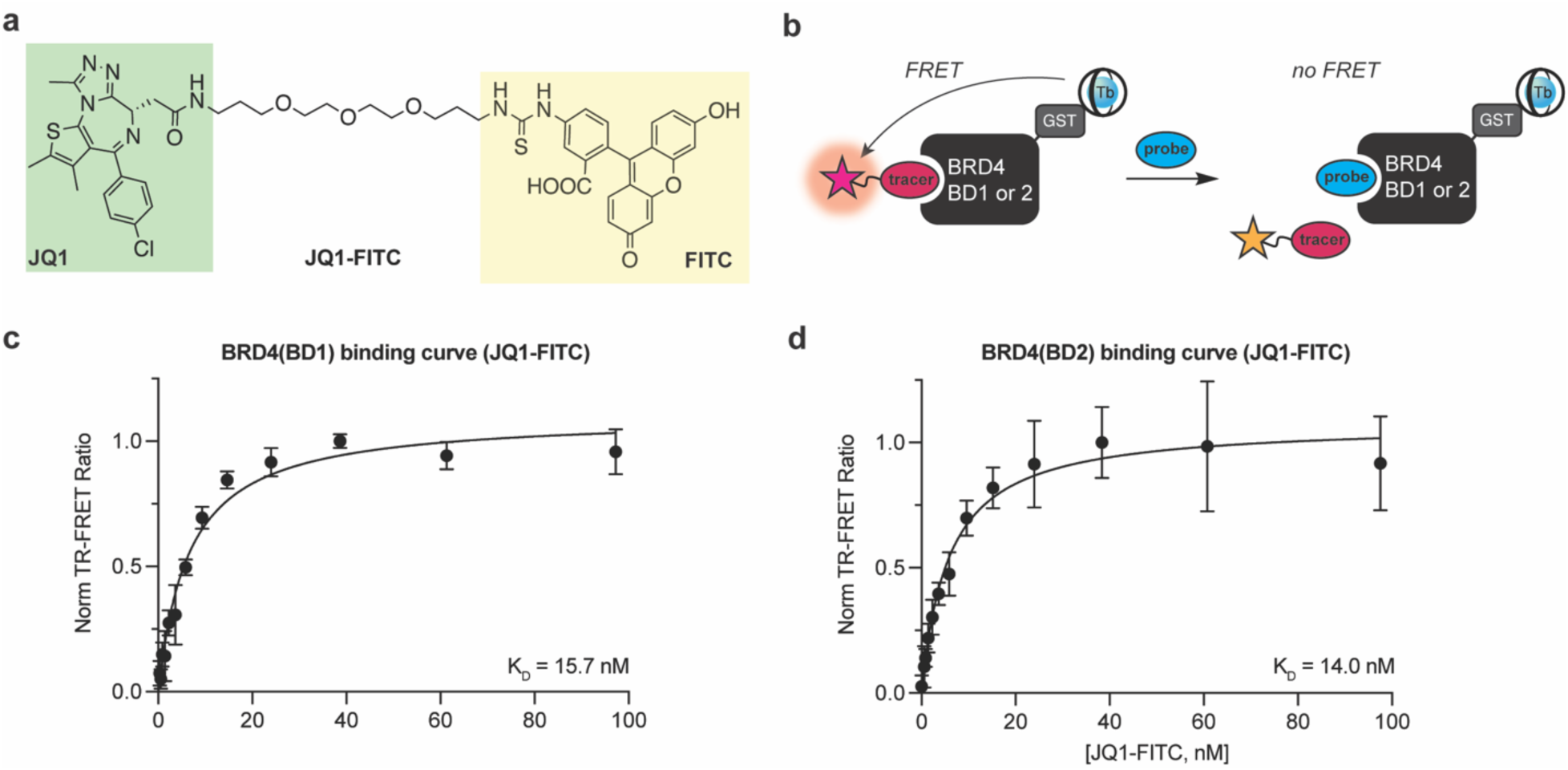
TR-FRET ligand displacement assay platform with GST-BRD4(BD1) and GST-BRD4(BD2). (**a**) Structure of JQ1-FITC, a tracer used to assess the engagement with BD1 and BD2 domains of BRD4. (**b**) Schematic of the TR-FRET ligand displacement assay using GST-BRD4(BD1) and GST-BRD4(BD2) with CoraFluor-1-labeled anti-GST nanobody. (**c**) Saturation binding of JQ1-FITC to GST-BRD4(BD1) labeled with CoraFluor-1-anti-GST nanobody. (**d**) Saturation binding of JQ1-FITC to GST-BRD4(BD2) labeled with CoraFluor-1-anti-GST nanobody. Data in (**c**) and (**d**) are presented as mean ± SD (n = 3 technical replicates).

**Extended Data Fig. 3.**
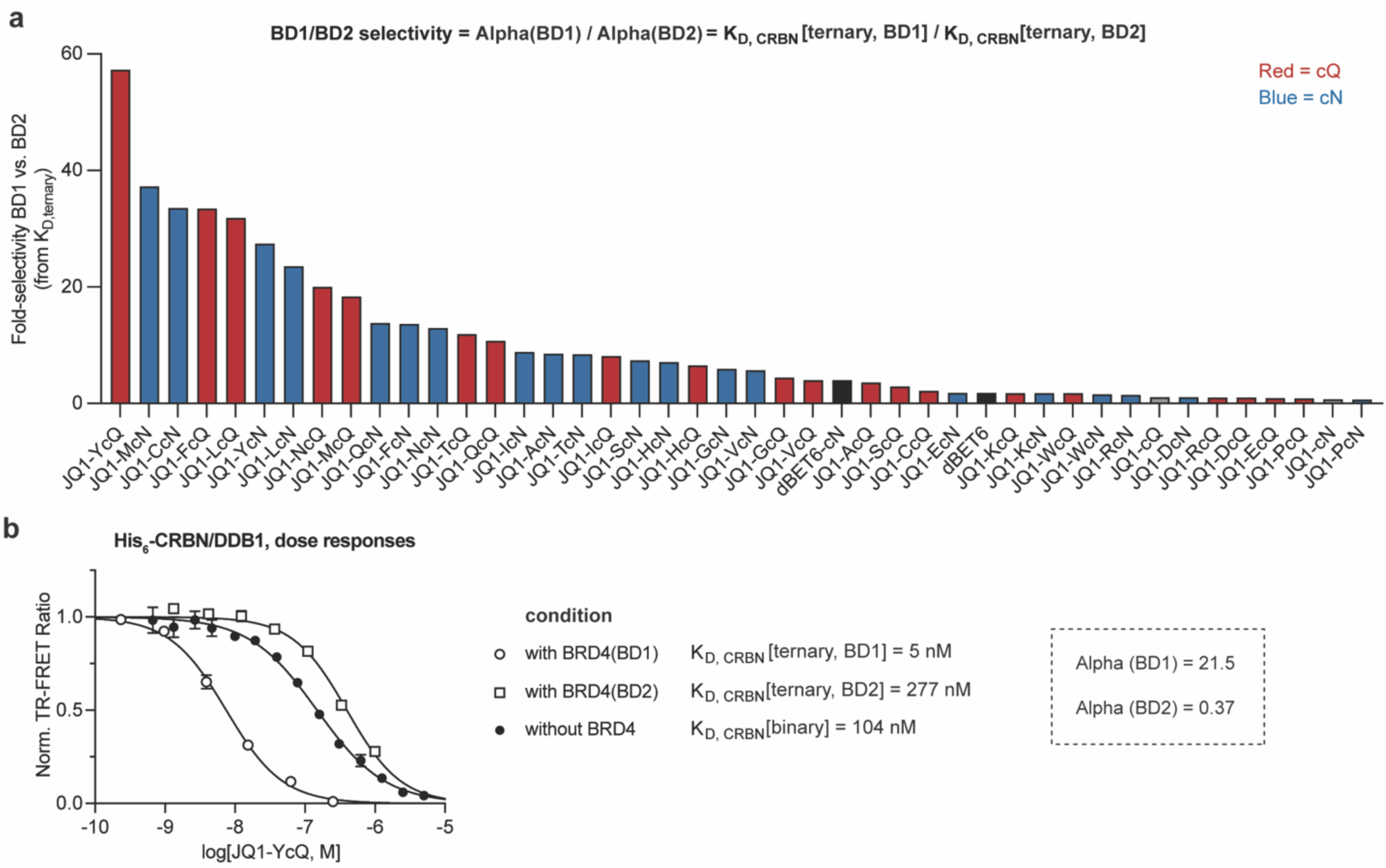
Evaluation of ternary complexes between CRBN/DDB1 and individual bromodomains of BRD4 (BD1 or BD2) induced by IMiD- and cyclimid-based BRD4 degraders. (**a**) Selectivity parameter between BD1 and BD2 degradation of IMiD-based degraders (black), cQ-cyclimid (red), and cN-cyclimid (blue) BRD4 degraders. Cyclimid degraders display higher biochemical selectivity than dBET6. (**b**) Dose-titration of JQ1-YcQ in TR-FRET ligand displacement assays with His_6_-CRBN/DDB1 in the absence of BRD4, or in the presence of a near-saturating concentration of either the BD1 or BD2 domain of BRD4 (1 µM). The obtained dissociation constants (K_D, CRBN_[binary] or K_D, CRBN_[ternary]) of JQ1-YcQ against the His_6_-CRBN/DDB1 complex indicated a 60-fold difference between K_D, CRBN_[ternary, BD1] and K_D, CRBN_[ternary, BD2], *i.e*., Alpha(BD1) and Alpha(BD2). Data are presented as mean ± SD (n = 3 technical replicates).

**Extended Data Fig. 4.**
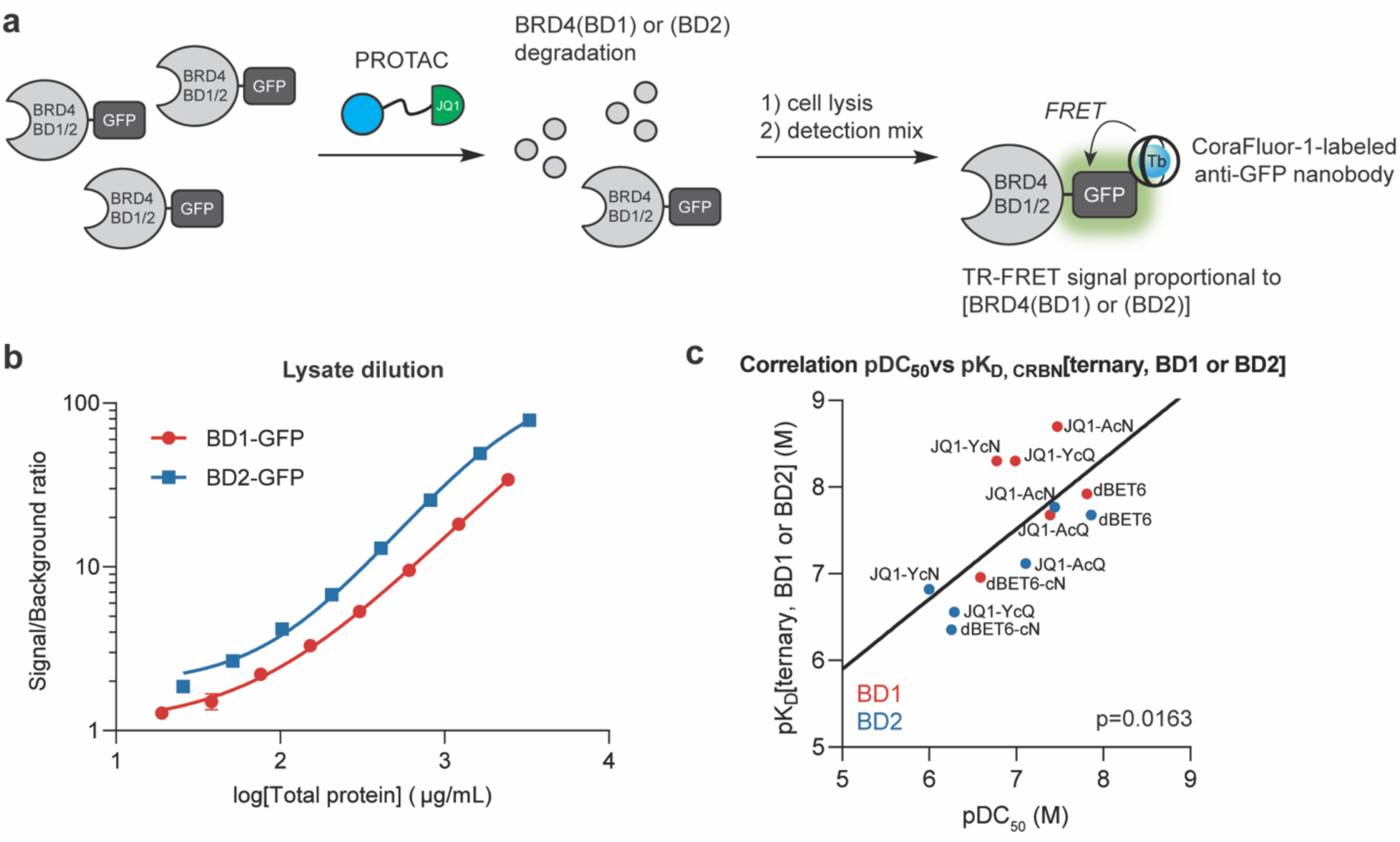
TR-FRET assay platform to quantitatively measure BRD4(BD1)-GFP and BRD4(BD2)-GFP levels in whole cell extracts. (**a**) Schematic showing the TR-FRET assay design for quantification of BRD4(BD1) and BRD4(BD2) levels in cell lysates. (**b**) TR-FRET-based quantification of BRD4(BD1)-GFP and BRD4(BD2)-GFP in serially diluted cell lysates from HEK293T cells stably expressing BRD4(BD1)-GFP or BRD4(BD2)-GFP. (n = 2 technical replicates, error bars, which fall within symbols, represent mean ± SD). (**c**) Correlation between pDC_50_ values (half-maximal degradation concentrations; DC_50_) of the selected cyclimid degraders for BRD4(BD1) or BRD4(BD2) degradation and their biochemical pK_D, CRBN_[ternary] values.

**Extended Data Fig. 5.**
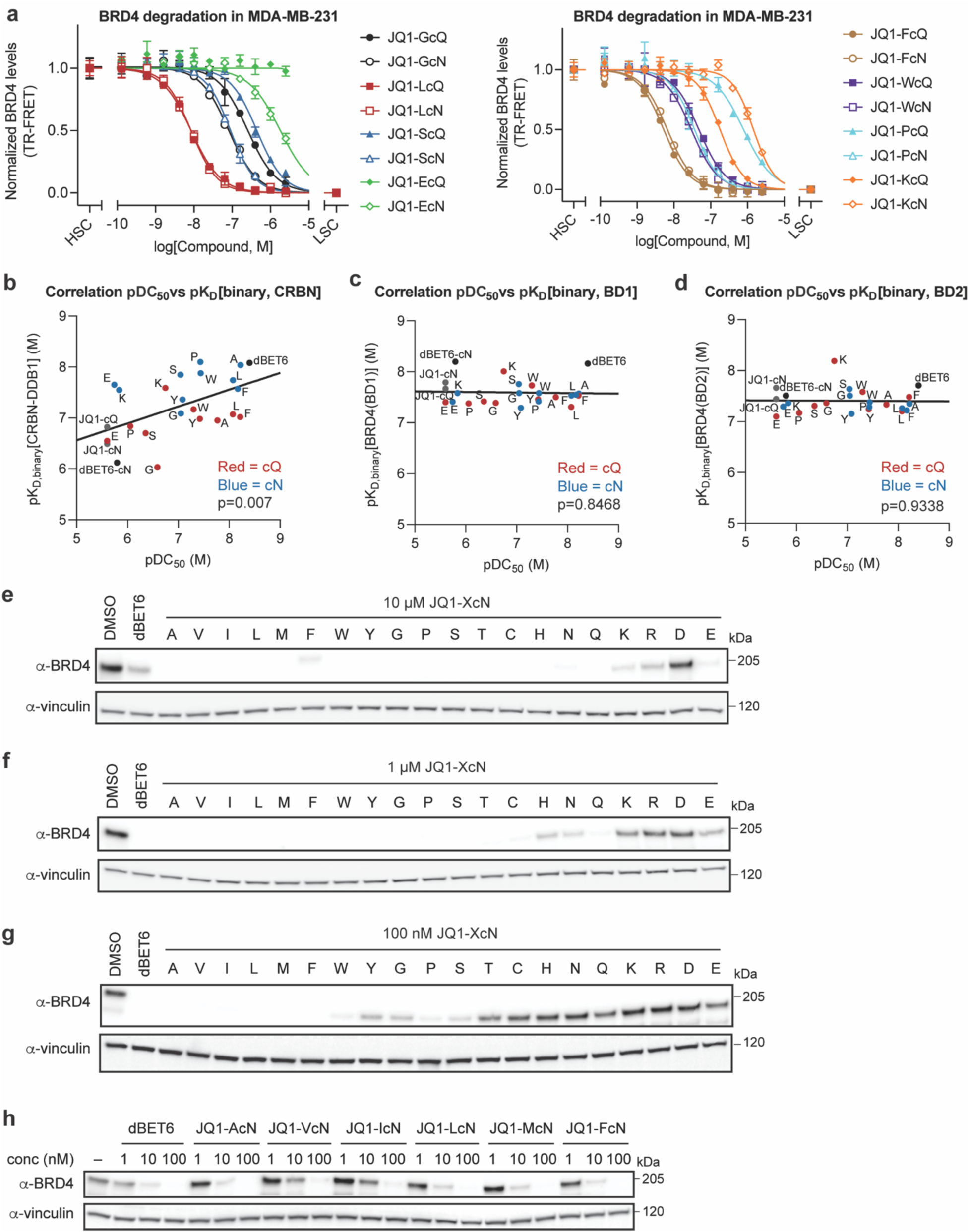
Cellular degradation assays of cyclimid-based BRD4 degraders. (**a**) BRD4 levels in MDA-MB-231 cell lysate after 5 h treatment with selected cyclimid BRD4 degraders were measured by TR-FRET assay. HSC, high signal control; LSC, low signal control. Data are presented as mean ± SD (n = 2 biologically independent samples). (**b–d**) Correlation between pDC_50_ values of the selected cyclimid degraders for endogenous BRD4 degradation and their pK_D_[binary] values against (**b**) CRBN, (**c**) BRD4(BD1), or (**d**) BRD4(BD2). (**e–g**) Western blots of BRD4 levels after treatment of HEK293T cells for 4 h with (**e**) 10 μM, (**f**) 1 μM, or (**g)** 100 nM dBET6 or the twenty cN-cyclimid BRD4 degraders. (**h**) Western blot of BRD4 levels after treatment of HEK293T cells with dBET6 and selected cN-cyclimid degraders for 4 h over a 1–100 nM dose-response range. All western blot data are representative of at least three independent replicates. For uncropped western blot images, see Supplementary Figure 3.

**Extended Data Fig. 6.**
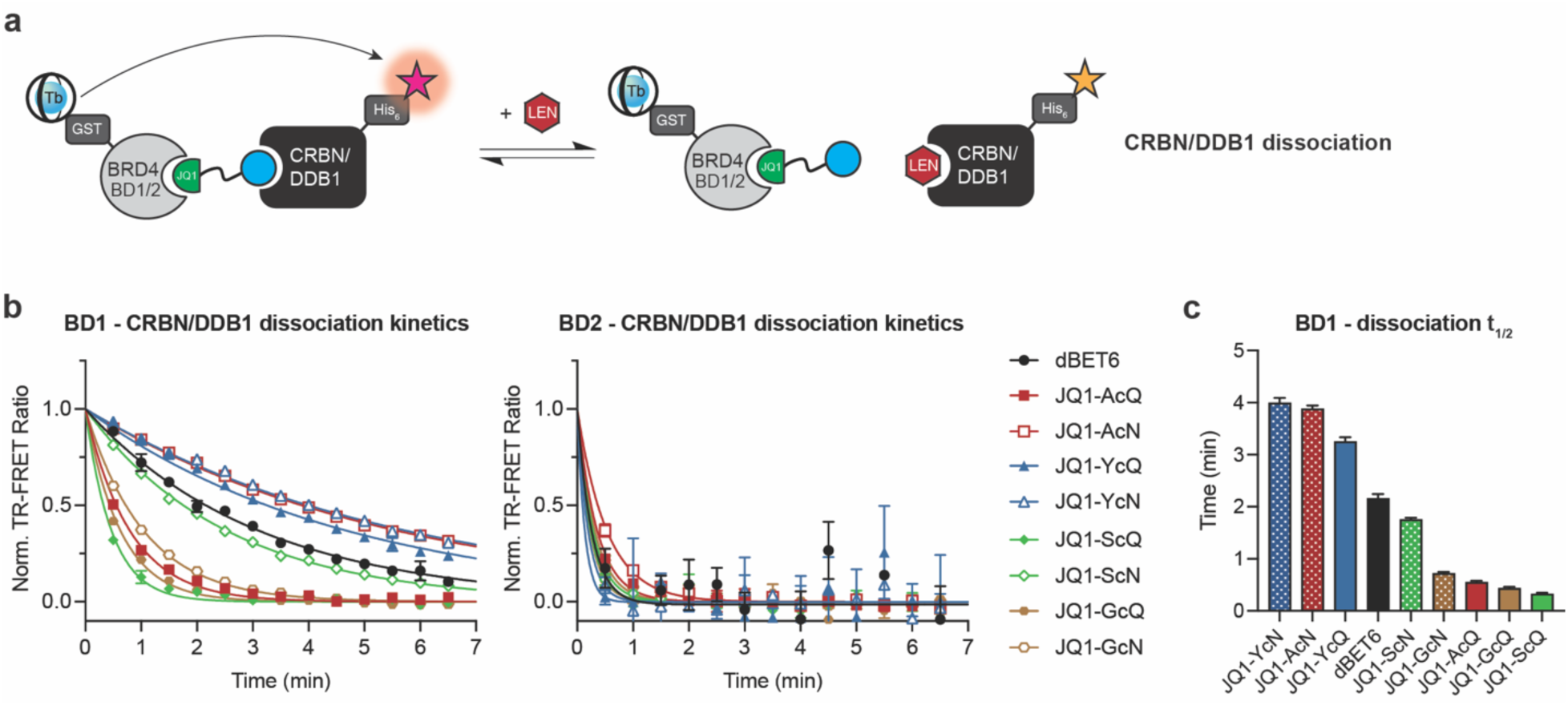
Ligand-dependent dynamics of CRBN/DDB1 dissociation from cyclimid degrader-induced ternary complexes with individual BRD4 bromodomains. (**a**) Schematic illustration of TR-FRET assay system employed to monitor CRBN/DDB1 dissociation from the degrader-mediated ternary complex. (**b**) Kinetic measurements of CRBN/DDB1 dissociation from the ternary complex induced by the selected cyclimids or dBET6. Data are presented as mean ± SD (n = 2 technical replicates) and fitted to a one-phase decay model in Prism 9. (**c**) Half-lives (*t*_1/2_) of the CRBN/DDB1–BRD4(BD1) ternary complex mediated by the indicated cyclimid degraders and dBET6. Error bars represent mean ± SD (n = 2 technical replicates).

**Extended Data Fig. 7.**
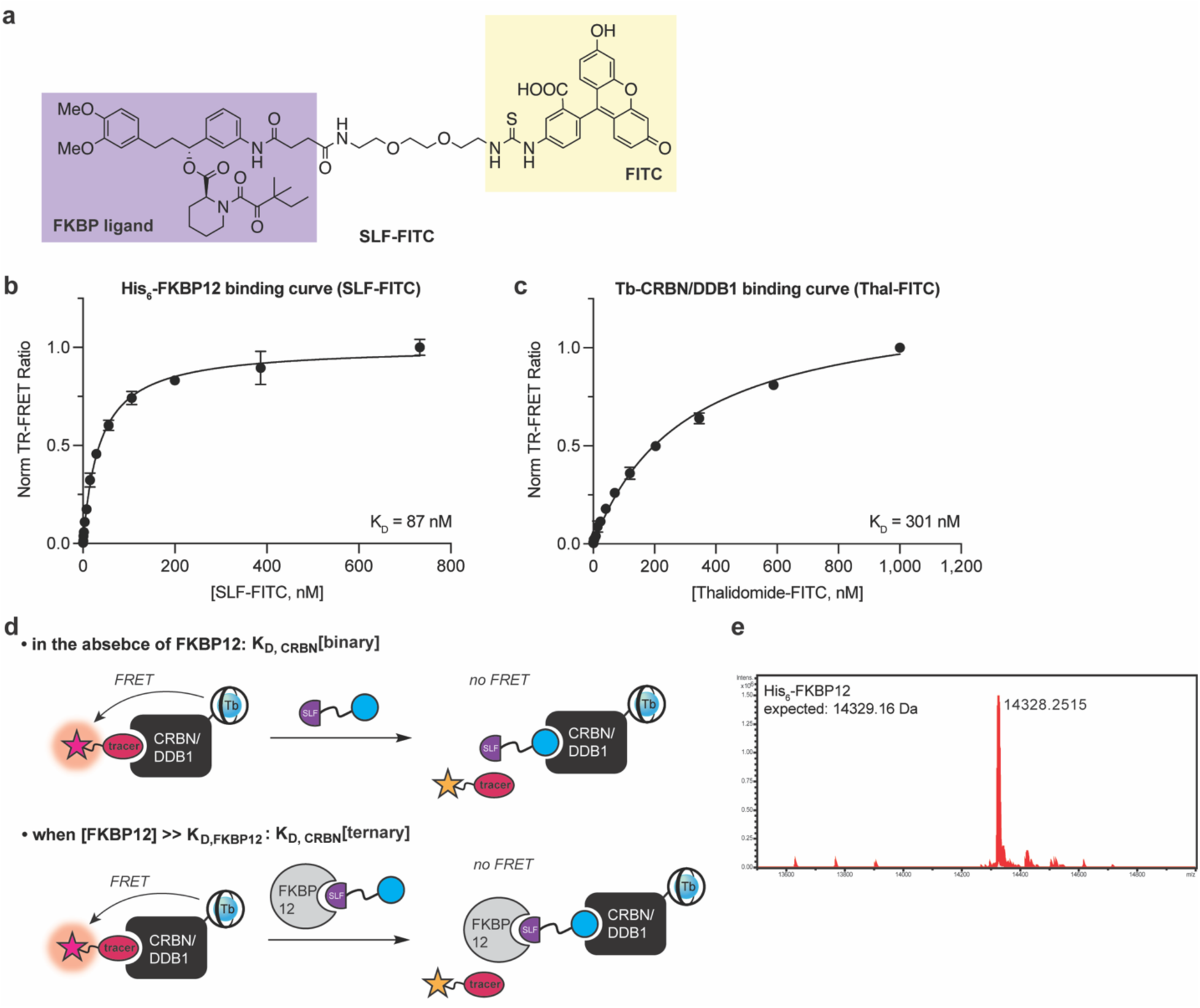
TR-FRET ligand displacement assay platform to measure small molecule target engagement with His_6_-FKBP12 and His_6_-CRBN/DDB1. (**a**) Structure of SLF-FITC, a tracer used to assess the engagement with FKBP12. (**b**) Saturation binding of SLF-FITC to His_6_-FKBP12 labeled with CoraFluor-1-anti-His_6_ conjugate. (**c**) Saturation binding of Thal-FITC to CoraFluor-1-labeled His_6_-CRBN/DDB1. Data in (**b**) and (**c**) are presented as mean ± SD (n = 3 technical replicates). (**d**) Schematic of the TR-FRET assay principle for determining K_D_[binary] and K_D_[ternary] against CRBN/DDB1, in the absence or the presence of FKBP12. (e) Intact MS measurement of His_6_-FKBP12.

**Extended Data Fig. 8.**
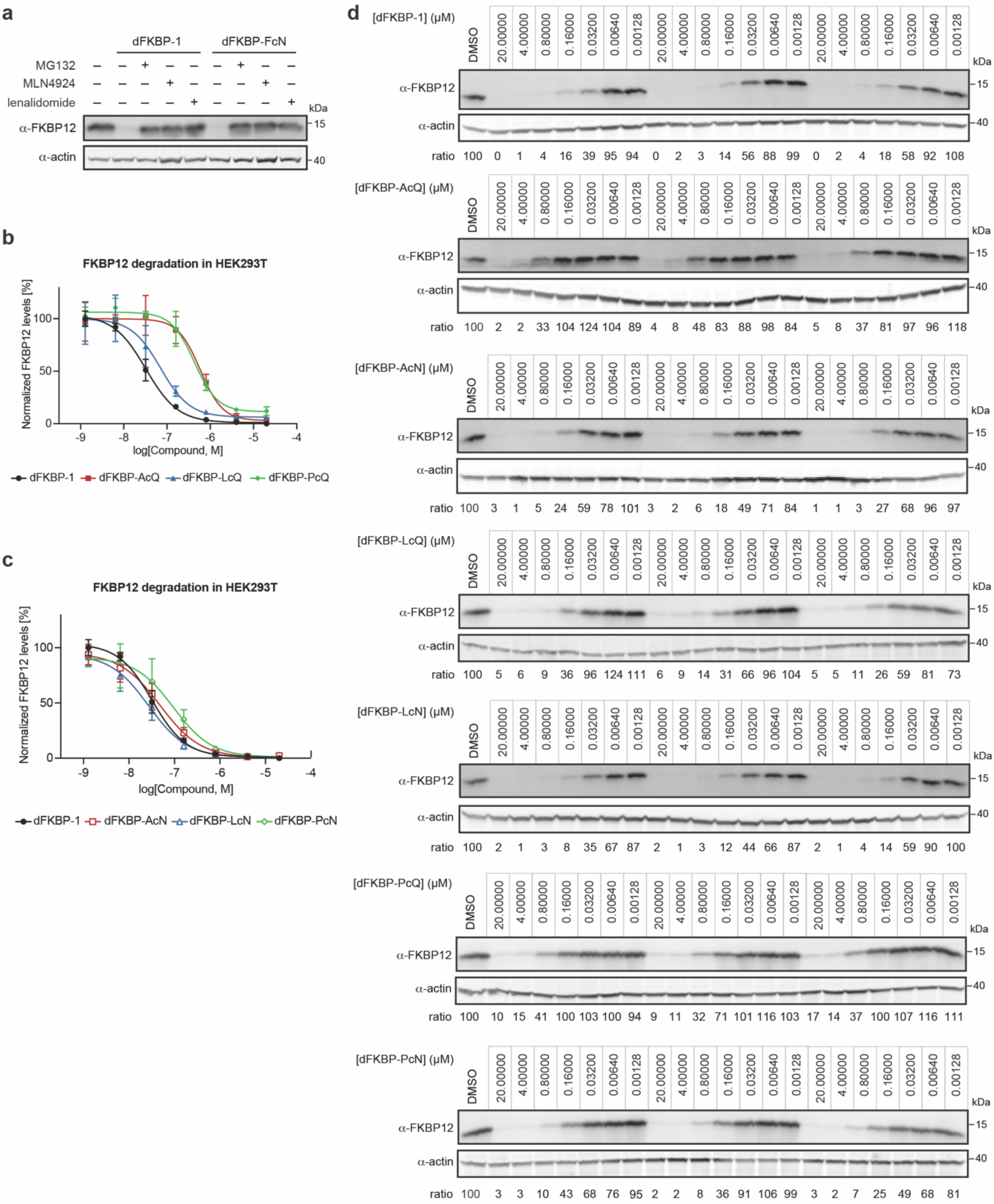
Evaluation of dFKBP-cyclimids in targeted protein degradation of FKBP12. (**a**) FKBP12 levels in HEK293T cells treated with the indicated degrader in the absence or the presence of a proteasome inhibitor MG132 (10 μM), a neddylation inhibitor MLN4924 (1 μM), or excess lenalidomide (100 μM), and incubated for 24 h. Rescue of FKBP12 degradation was observed when cells were pre-treated with MG132, MLN4924, or lenalidomide. (**b–c**) FKBP12 levels in HEK293T cells treated with serially diluted degraders for 24 h. Quantification of FKBP12 levels were calculated relative to DMSO control, and the plots shown were used to calculate the tabulated half-maximal degradation concentration (DC_50_) values for each degrader. Data in (**b**) and (**c**) are presented as mean ± SD (n = 3 biologically independent samples). (**d**) Corresponding western blots of FKBP12 levels after treatment of HEK293T cells for 24 h with dFKBP-1 or each of the selected dFKBP-cyclimids over a dose-response range of 1.28 nM and 20 μM. All western blot data are representative of at least three independent replicates. For uncropped western blot images, see Supplementary Figure 4.

**Extended Data Fig. 9.**
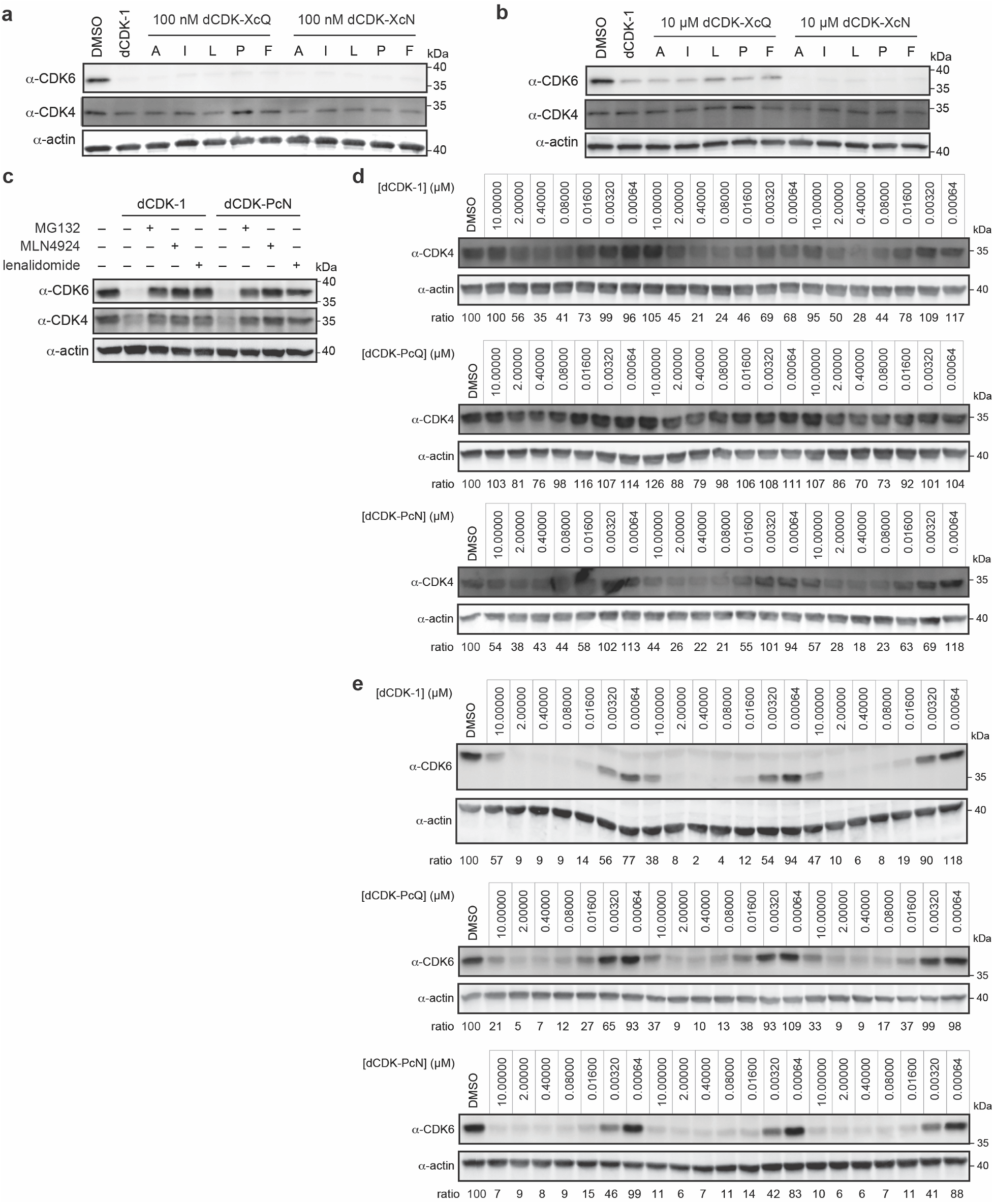
Evaluation of dCDK-cyclimids in targeted protein degradation of CDK4 and CDK6. (**a**) Western blot of CDK4 and CDK6 levels after treatment of Jurkat cells for 4 h with 100 nM dCDK-1 or each dCDK-cyclimids. (**b**) Western blot of CDK4 and CDK6 levels after treatment of Jurkat cells for 4 h with 10 μM dCDK-1 or each dCDK-cyclimids. (**c**) CDK4 and CDK6 levels in Jurkat cells treated with the indicated degrader in the absence or the presence of a proteasome inhibitor MG132 (10 μM), a neddylation inhibitor MLN4924 (1 μM), or excess lenalidomide (100 μM), and incubated for 4 h. Rescue of CDK4 and CDK6 degradation was observed when cells were pre-treated with MG132, MLN4924, or lenalidomide. (**d–e**) Corresponding western blots of (**d**) CDK4 and (**e**) CDK6 after treatment for 4 h with dCDK-1, dCDK-PcQ, or dCDK-PcN over a dose-response range of 0.64 nM and 10 μM. Quantification of CDK4 and CDK6 levels were calculated relative to DMSO control to obtain the plots in Figure 5h–j. All western blot data are representative of at least three independent replicates. For uncropped western blot images, see Supplementary Figure 5.

**Extended Data Fig. 10.**
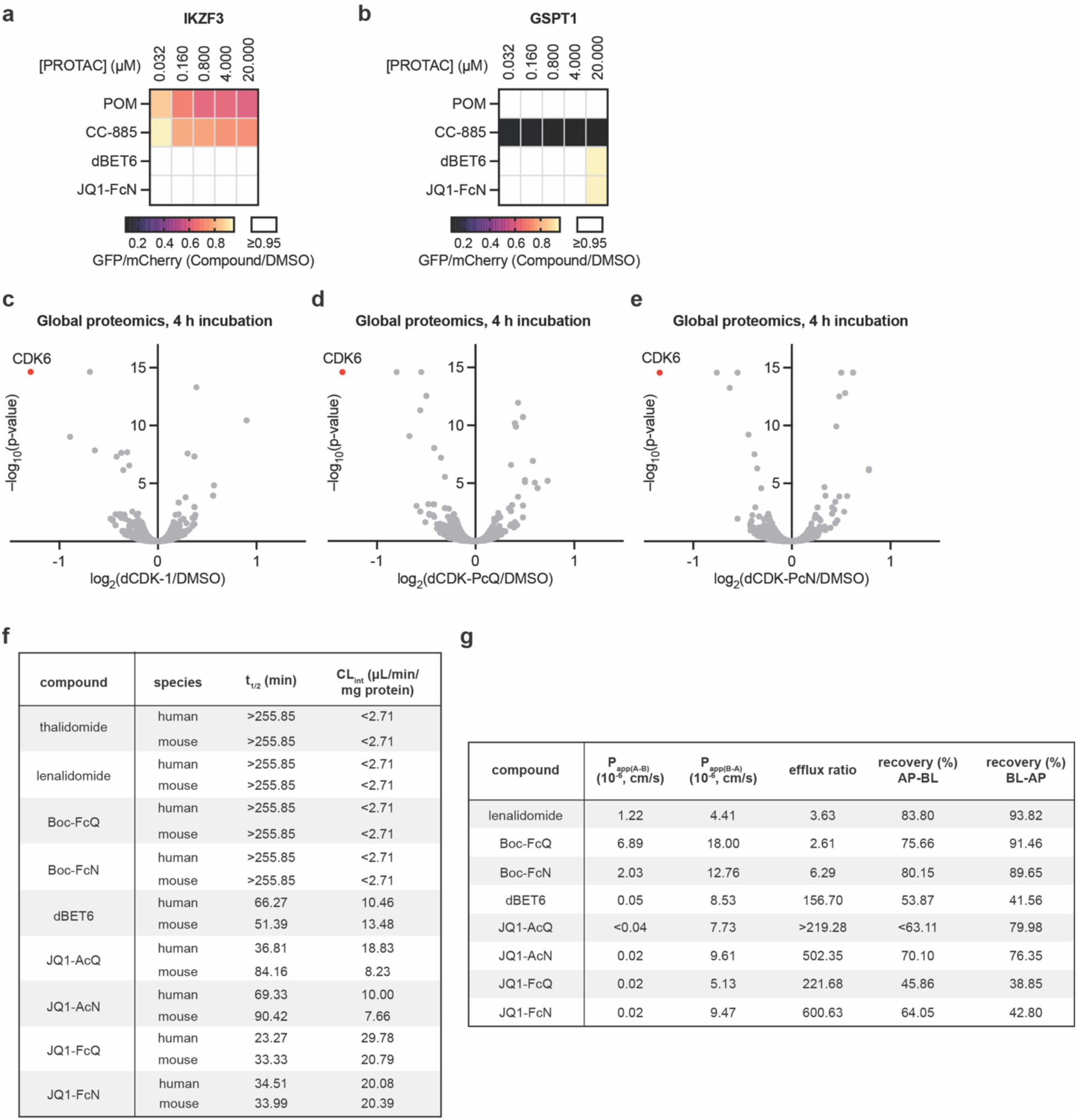
The cyclimids have minimal IMiD-sensitive off-target degradation and possess comparable metabolic stability and cell permeability properties to the IMiDs. (**a**) Degradation of IKZF3 ZF degrons in cells by pomalidomide, CC-885, dBET6, and JQ1-FcN over a dose-response range of 32 nM to 20 μM. U2OS cells stably expressing IKZF3 ZF degrons fused to GFP were treated with degraders followed by flow cytometry to assess IKZF3 ZF degradation. (**b**) Degradation of GSPT1 in cells using pomalidomide, CC-885, dBET6, and JQ1-FcN. HEK293T cells stably expressing GSPT1-GFP were treated with the indicated compounds followed by flow cytometry to assess GSPT1 degradation. Data in (**a**)–(**b**) are presented as mean (n = 3 biologically independent samples). (**c–e**) Quantitative proteomics of Jurkat cells after 4 h treatment with 0.1 μM of (**c**) dCDK-1, (**d**) dCDK-PcQ, or (**e**) dCDK-PcN. P-values for the abundance ratios were calculated by one-way ANOVA with TukeyHSD post-hoc test. (**f**) Metabolic stability of the selected monofunctional cyclimids, thalidomide, and lenalidomide, as well as heterobifunctional degraders bearing the cyclimids or the IMiDs (dBET6), against liver S9 fractions. (**g**) *in vitro* permeability of the selected monofunctional cyclimids and lenalidomide, as well as heterobifunctional degraders bearing the cyclimids or the IMiDs (dBET6), in a Caco-2 trans-well assay.

